# Apremilast improves cardiomyocyte cohesion and arrhythmia in different models for arrhythmogenic cardiomyopathy

**DOI:** 10.1101/2025.04.17.649297

**Authors:** Konstanze Stangner, Orsela Dervishi, Janina Kuhnert, Carl Wendt, Soumyata Pathak, Maria Shoykhet, Silvana Olivares Florez, Sina Moztarzadeh, Jens Opsteen, Ni Luh Cathrin Suniasih Wohlfarth, Ruth Biller, Elisabeth Graf, Dominik S. Westphal, Tatjana Williams, Brenda Gerull, Tomo Šarić, Sunil Yeruva, Jens Waschke

**Affiliations:** Chair of Vegetative Anatomy, Institute of Anatomy, Faculty of Medicine, Ludwig-Maximilian-University (LMU) Munich, 80336 Munich, Germany; University of Cologne, Faculty of Medicine and University Hospital Cologne, Center for Physiology and Pathophysiology, Institute for Neurophysiology, Cologne, Germany; ARVC-Selbsthilfe e.V., patient association, Munich, Germany; Institute of Human Genetics, School of Medicine, Technical University of Munich, Munich, Germany; Department of Internal Medicine I, Technical University of Munich and University Hospital, School of Medicine and Health, Technical University of Munich, Munich, Germany; Comprehensive Heart Failure Center and Department of Medicine I, University Hospital Würzburg, Würzburg, Germany

**Author notes:** Department of Otorhinolaryngology, Technical University of Munich and University Hospital, Ismaningerstrasse 22, 81675, Munich, Germany (Maria Shoykhet). These authors contributed equally (Konstanze Stangner, Orsela Dervishi, Tomo Šarić, Sunil Yeruva, Jens Waschke). **Corresponding authors:** Tomo Šarić, Institute for Neurophysiology, Faculty of Medicine and University Hospital Cologne, University of Cologne Robert Koch Str. 39, 50931 Cologne Germany, Sunil Yeruva and Jens Waschke, Chair of Vegetative Anatomy, Institute of Anatomy, Ludwig-Maximilians-University, Pettenkoferstrasse 11, 80336 Munich, Germany; and.

**Keywords:** Apremilast, arrhythmogenic cardiomyopathy, arrhythmia, desmoplakin and desmoglein 2

## Abstract

**Background:** Arrhythmogenic cardiomyopathy (ACM) is a genetically inherited desmosome heart disease leading to life-threatening arrhythmias and sudden cardiac death. Currently, ACM treatment paradigms are merely symptom targeting. Recently, apremilast was shown to stabilize keratinocyte adhesion in the desmosomal disease pemphigus vulgaris. Therefore, this study investigated whether apremilast can be a therapeutic option for ACM.

**Methods:** Human induced pluripotent stem cells from a healthy control (hiPSC) and an ACM index patient (ACM-hiPSC) carrying a heterozygous desmoplakin (*DSP*) gene mutation (c.2854G>T, p.Glu952Ter), confirmed by whole exome sequencing (WES), were established. Cyclic-AMP ELISA, dissociation assay, immunostaining, and Western blotting analyses were performed in human iPSC-derived cardiomyocytes (hiPSC-CMs), murine HL-1 cardiomyocytes, and cardiac slices derived from wild-type (WT) mice, plakoglobin (PG, *Jup*) knockout (*Jup^-/-^*) (murine ACM model) or PG Serine 665 phosphodeficient (JUP-S665A) mice. Microelectrode array (MEA) analyses in ventricular cardiac slices and Langendorff heart perfusion were performed to analyze heart rate variability and arrhythmia.

**Results:** ACM-hiPSC derived cardiomyocytes (ACM-hiPSC-CMs) revealed a significant loss of cohesion, which was rescued by apremilast. Further, treatment with apremilast strengthened basal cardiomyocyte cohesion in HL-1 cells and WT murine cardiac slices, paralleled by phosphorylation of PG at Serine 665 in human and murine models. In HL-1 cells, apremilast in addition activated ERK1/2, inhibition of which abolished apremilast-enhanced cardiomyocyte cohesion. Further, dissociation assays in slice cultures from JUP-S665A and *Jup^-/-^* mice revealed that PG is crucial for apremilast-enhanced cardiomyocyte cohesion. In parallel to enhanced cell adhesion, MEA and Langendorff measurements from WT and *Jup^-/-^* mice demonstrated decreased heart rate variability and arrhythmia after apremilast treatment.

**Conclusions:** Apremilast improves loss of cardiomyocyte cohesion, enhances localization of DSG2, and reduces arrhythmia in human and murine models of ACM *ex vivo* and *in vitro*, providing a novel treatment strategy for ACM by preserving desmosome function.

**Translational perspective:** The current therapeutic options for patients with arrhythmogenic cardiomyopathy (ACM) include lifestyle changes, treatment with anti-arrhythmic drugs, catheter ablation, implantable cardiac defibrillators, and ultimately, heart transplantation for patients who are having therapy refractory arrhythmia or developed heart failure. However, lifestyle changes, such as restraining from physical endurance activities and β-blocker therapy, are most used in patients carrying genetic variants coding for proteins of the desmosomal complex. Recent advancements hint that strategies enhancing intracellular cAMP could be beneficial in treating desmosomal diseases and can be effective therapeutics, which would be highly relevant for ACM patients.

In this study, we show that apremilast improves loss of cardiomyocyte cohesion, enhances localization of desmosomal proteins, and reduces arrhythmia in both human and murine models of ACM ex vivo and in vitro, providing a novel treatment strategy for ACM by preserving desmosome function.

## Introduction

Arrhythmogenic cardiomyopathy (ACM) is an uncommon hereditary heart condition that can lead to sudden cardiac death (SCD), particularly among young athletes ^1, 2^. The estimated prevalence of this condition ranges from 1 in 1000 to 1 in 5000 individuals ^3, 4^. Notably, SCD can even occur in the initial concealed phase without observable structural changes in the ventricles, and it may be the first manifestation of the disease in approximately 5-10% of cases ^3–5^.

Approximately 60% of ACM patients carry pathogenic variants in genes coding for desmosomal proteins of the intercalated disc (ICD) ^6, 7^. ICDs serve as specialized intercellular contacts where desmosomes and adherens junctions (AJ), which provide mechanical adhesion, together with gap junctions (GJ) and sodium channels crucial for electric coupling, form a complex structure known as the connexome ^8–10^. Within the connexome, the propagation of excitation is facilitated by GJ through electrotonic and ephaptic coupling with Nav1.5 sodium channels.

In individuals with ACM, pathogenic variants in genes encoding desmosomal proteins encompass both plaque proteins like plakophilin 2 (*PKP2*), desmoplakin (*DSP*), and plakoglobin (*JUP*), as well as cadherin-type adhesion molecules such as desmoglein 2 (*DSG2*) and desmocollin 2 (*DSC2)* ^11–13^. Pathogenic variants affecting N-cadherin (*CDH2*), desmin (*DES*), transmembrane protein 43 (*TMEM43*), phospholamban (*PLN*), filamin-C (*FLNC*) and sodium voltage-gated channel subunit 5 (*SCN5A* or Nav1.5) are rarely detected ^14–17^. Hence, ACM is recognized primarily as a hereditary condition affecting the desmosome. Pathogenic variants in desmosome genes such as *DSG2, DSP,* and *PKP2* associated with ACM often disrupt intercellular desmosomal connections, thereby reducing cellular cohesion^8^. Recently, it has been shown that pathogenic variants in desmosomal genes are not confined to ACM but are present in myocarditis patients, especially pathogenic variants in *DSP* ^18–20^. Moreover, in the last couple of years, autoantibodies against ICD proteins have been detected in ACM patients, which were shown to reduce cardiomyocyte cohesion by inhibition of DSG2 and N-cadherin (N-CAD) binding ^21–23^.

Various pathways regulating cardiomyocyte cohesion have been identified ^8, 24–27^. Additionally, mounting evidence suggests that desmosomes play a role in cellular cohesion and can serve as signaling centers and maintain tissue integrity ^8, 28–31^. Thus, cardiac desmosomes are integral components of an intricate signaling network that can both influence and be influenced by cellular signaling processes.

The current therapeutic options for patients with ACM include lifestyle changes, treatment with anti-arrhythmic drugs, catheter ablation, implantable cardiac defibrillators, and ultimately, heart transplantation for patients who are having therapy refractory arrhythmia or developed heart failure ^1, 32^. However, in patients carrying genetic variant coding for proteins of the desmosomal complex, lifestyle changes, such as restraining from physical endurance activities and β-blocker therapy, are most commonly used ^1, 33, 34^.

In our previous work, we discovered increased cAMP levels via PKA-mediated PG phosphorylation at serine 665 (PG-pS665) as a driving force for DSG2-mediated enhanced cardiomyocyte cohesion, which is referred to as positive adhesiotropy ^8, 24, 25^. This indicated that cardiomyocyte adhesion at ICDs is precisely regulated and can be enhanced via cAMP-increasing drugs. In addition to the above evidence, recent advances in research suggest that similar molecular mechanisms are observed in ACM and pemphigus vulgaris. Pemphigus is a desmosomal disease of the skin similar to ACM but caused by autoantibodies primarily targeting desmosomal cadherins ^8, 30^. We have recently demonstrated that apremilast, a PDE-4 inhibitor, clinically used in treating psoriasis and Behçet’s disease, abrogated pemphigus autoantibody-induced loss of keratinocyte cohesion via PG-pS665, inhibition of autoantibody-mediated keratin retraction and desmoplakin (DP) assembly in the desmosomal plaques ^35^. Further, two recent case report studies showed that apremilast treatment in a pemphigus patient led to a decrease in DSG-specific autoantibody levels and the healing of lesions ^36, 37^.

These recent advancements hint that strategies enhancing intracellular cAMP could be beneficial in treating desmosomal diseases and can be effective therapeutics, which would be highly relevant for ACM patients. Therefore, this study investigated whether apremilast can enhance desmosomal-mediated cardiomyocyte cohesion and be used as an ACM therapeutic agent.

## Results

### Apremilast enhances cardiomyocyte cohesion, DSG2-DP translocation, PG phosphorylation and ERK1/2 activation in HL-1 cells

First, we measured cAMP production in HL-1 cells after treatment for 30 min with the combination of 5 µM forskolin and 10 µM rolipram (FR), as well as 1 µM and 10 µM apremilast. We observed an increase in cAMP from 14.02±4.02 pmol/ml in control to 334.1±83.72, 24.68±4.40 and 27.43±5.11 pmol/ml after treatment with FR, 1 µM apremilast, and 10 µM apremilast, respectively (Fig. 1a). Further, we analyzed HL-1 cardiomyocyte cohesion utilizing dissociation assays. Treatment with FR and apremilast at both concentrations (1 µM and 10 µM) led to a significant increase in basal cell cohesion as indicated by a lower number of monolayer fragments (Fig. 1b). Since 1 µM apremilast corresponds to the plasma concentration of apremilast administered to humans ^35^, we continued using this concentration further. In addition to the HL-1 cells, we also performed dissociation assays in WT murine cardiac slice cultures and found that similar to HL-1 cells, apremilast enhanced cardiomyocyte cohesion in intact cardiac tissue (Fig. 1c).

**Figure 1:**
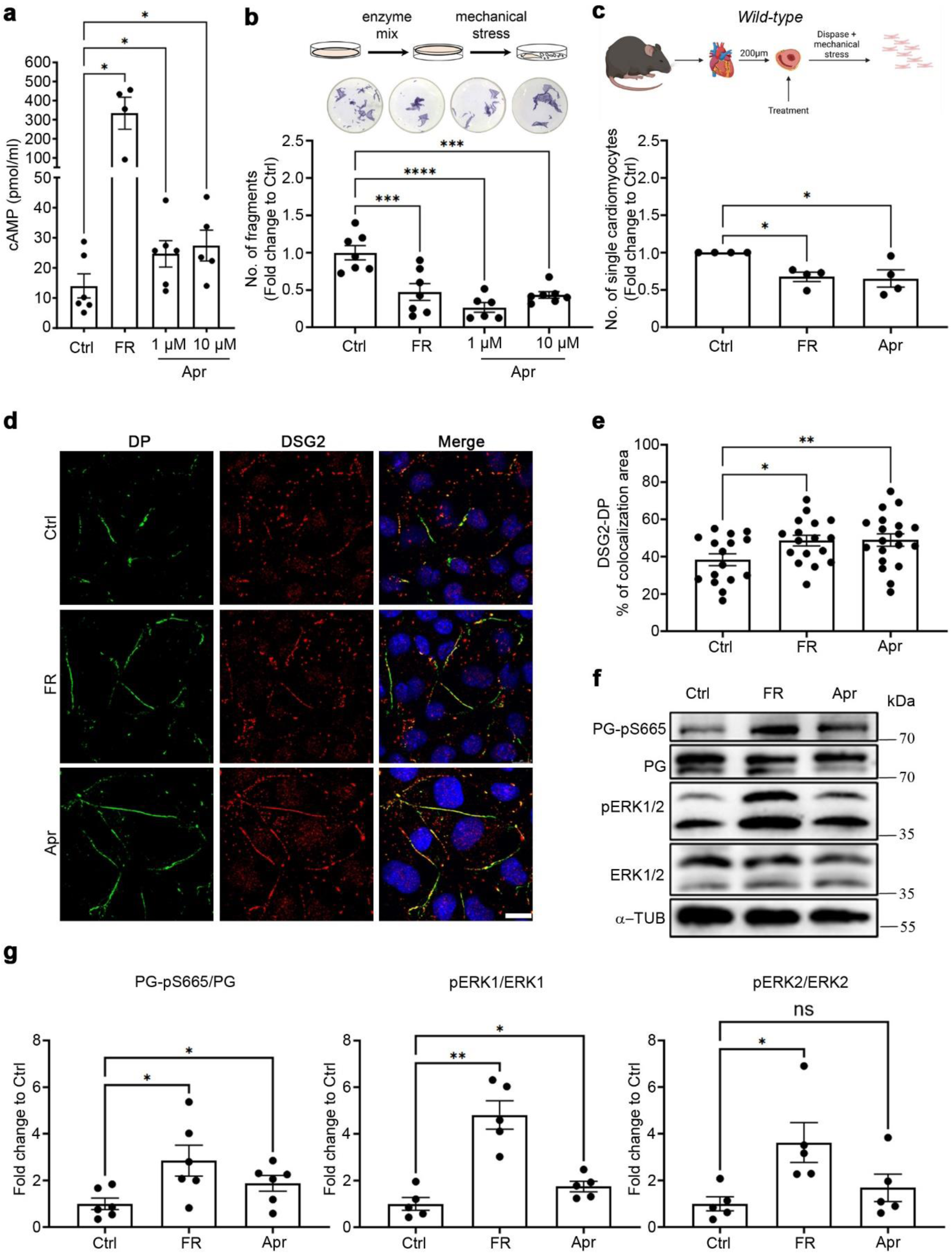
Apremilast (Apr) enhances cardiomyocyte cohesion in HL-1 cells and wild-type murine cardiac slices. **a.** HL-1 cells were treated with apremilast (Apr, 1 and 10 µM) for 30 min, and cAMP was measured; FR was used as a positive control in all assays. N=4-6. **b.** Dissociation assays were performed in HL-1 cells treated with 1 or 10 µM Apr for 1 h. N=7. The cartoon represents the dissociation assay method, and the panel below displays representative wells for each condition. MTT was used to show the viability of cells. **c.** Dissociation assays were performed in wild-type murine cardiac slices treated with 1 µM Apr for 1 h. N=4. The cartoon represents the dissociation assay method. **d.** Immunostaining and confocal microscopy were performed for DP (green) and DSG2 (red) in HL-1 cells treated with FR and 1 µM Apr for 1 h. Scale bar: 10 µm. **e.** Quantification of DSG2-DP colocalization area. Each data point in the graph represents one confocal image performed across 3-4 experimental repeats. **f.** Representative Western blots for PG-pS665, PG, pERK1/2, and ERK1/2 in HL-1 cells treated with FR and 1 µM Apr for 1 h. α-tubulin was used as a loading control. **g.** Quantification of Western blots from **f.** N=5-6. All bar graphs in the figure represent mean±SEM. One-way ANOVA with Holm-Šidák post-hoc analysis was performed to assess the statistical significance. *p<0.05, **p<0.005, ***p<0.0005 and ****p<0.00005.

As we have previously established that enhanced intracellular cAMP leads to enhanced DP and DSG2 translocation, PG-pS665 and ERK1/2 activation ^24, 25, 27^, we investigated the same after treating HL-1 cells with apremilast and FR as a positive control. Immunostaining revealed that apremilast increased the translocation of DSG2 and DP to cell junctions (Fig. 1d and e). Western blot analysis showed that apremilast, similar to FR, enhanced phosphorylation of both PG at serine 665 and ERK1/2 (Fig. 1f and g).

### Apremilast-enhanced cardiomyocyte cohesion requires PG

We recently established that apremilast rescues impaired keratinocyte adhesion by phosphorylating PG at serine 665 ^35^. Therefore, we further investigated the role of PG serine 665 phosphorylation in cAMP-driven cardiomyocyte cohesion enhancement. For this, we utilized PG serine 665 phospho-deficient mice (JUPS665A)^35^. Hearts from these mice looked normal both morphologically and histologically (Suppl. Fig. 1a). A dissociation assay performed in murine cardiac slices from JUPS665A mice revealed enhanced cardiomyocyte cohesion after apremilast (Fig. 2a), revealing that apremilast might work through different signaling mechanisms in the absence of PG S665. We further investigated whether apremilast can enhance cardiomyocyte cohesion in a murine ACM model for which we utilized cardiac-specific PG knockout (*Jup^-/-^*) mice, which exhibited loss of cardiomyocytes and fibrosis similar to that observed in ACM patients ^24, 27^ (Suppl. Fig. 1b). Treatment of ventricular cardiac slices with apremilast did not enhance cardiomyocyte cohesion in *JUP^-/-^* mice (Fig. 2b). In contrast, in *Pkp2^-/-^*hearts derived from a murine ACM model ^38^, both apremilast and FR enhanced cardiomyocyte cohesion (Suppl. Fig. 2). We performed Western blot analysis to check for PG-pS665, which revealed that both apremilast and FR induced phosphorylation of PG at serine 665 in the wild-type JUPS665 mice. However, the PG-pS665 was absent in the JUPS665A mice (Fig. 2c). We then asked whether apremilast would enhance DSG2 translocation to strengthen the ICD. To answer this, we performed immunostaining in ventricular cardiac slices from WT, JUPS665A and *Jup^-/-^* mice and measured the thickness of staining for PG and DSG2 in the ICD, as described previously ^38^. Apremilast increased the thickness of PG and DSG2 staining in both WT (Fig. 2d and e) and JUPS665A (Fig. 2f) mice cardiomyocyte ICDs, whereas we did not detect either PG or DSG2 in *Jup^-/-^* mice cardiomyocyte ICDs, as observed in our previous studies^24, 25, 38, 39^, and apremilast did not further enhance either PG or DSG2.

**Figure 2:**
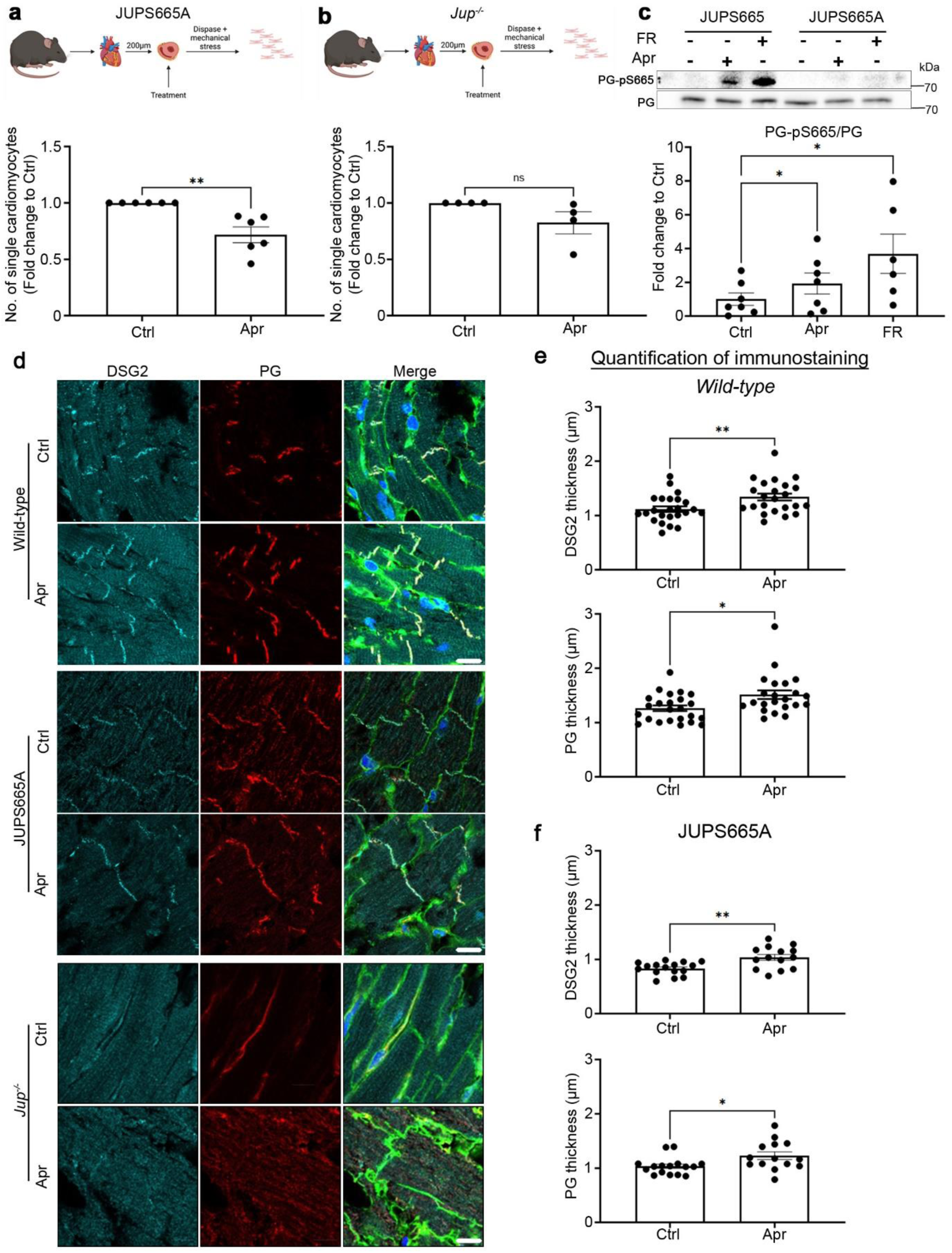
Apremilast requires plakoglobin to enhance cardiomyocyte cohesion. **a.** Dissociation assays were performed with cardiac slices of PG-S665 phosphorylation deficient mice (JUPS665A), N=5-6, and **b.** cardiomyocyte-specific deletion of PG (*Jup^-/-^*) mice treated with apremilast (Apr) for 1 h, N=4. The cartoons represent the dissociation assay method. **c.** Representative Western blots for PG-pS665 and PG, in cardiac slices from JUPS665 (WT) and JupS665A mice treated with Apr and FR for 1 h. The quantification of Western blots is displayed in the panel below. N=6-7. **d.** Immunostaining and confocal microscopy were performed for DSG2 (green) and PG (red) in cardiac slices from wild-type, JUPS665A and *Jup^-/-^* treated with Apr for 1 h. Scale bar 10 µm. **e.** The thickness of DSG2 and PG staining in the ICD was measured and represented in µm. Each data point in the graph represents the mean of multiple ICDs in one confocal image and measured across cardiac slices from 4-5 mice for each condition. Adjacent slices from a single mouse heart were used as a control for the respective treatments. All bar graphs in the figure represent mean±SEM. Paired student t-test (**b, e,** and **f**) or One-way ANOVA with Holm-Šidák post-hoc analysis (**a** and **c**) was performed to determine the statistical significance. *p<0.05 and **p<0.005.

### Activation of ERK1/2 is crucial in apremilast-enhanced cardiomyocyte cohesion

Furthermore, we investigated the role of ERK1/2 in apremilast-enhanced cardiomyocyte cohesion, as we found earlier that the positive effects of cAMP to stabilize cardiomyocyte cohesion were dependent on ERK1/2 ^25^. Here, we utilized HL-1 cells and treated them with apremilast alone or in combination with U0126, a MEK inhibitor upstream of ERK1/2. Dissociation assays revealed that after inhibition of ERK1/2 activation by U0126, apremilast did not enhance cardiomyocyte cohesion (Fig. 3a), as observed with FR in our previous study ^25^. Under these conditions, PG-pS665 was not altered by apremilast (Fig. 3b and c). However, apremilast-mediated DSG2 and DP translocation to cell borders was abolished in the absence of ERK1/2 activation (Fig. 3d and e).

**Figure 3:**
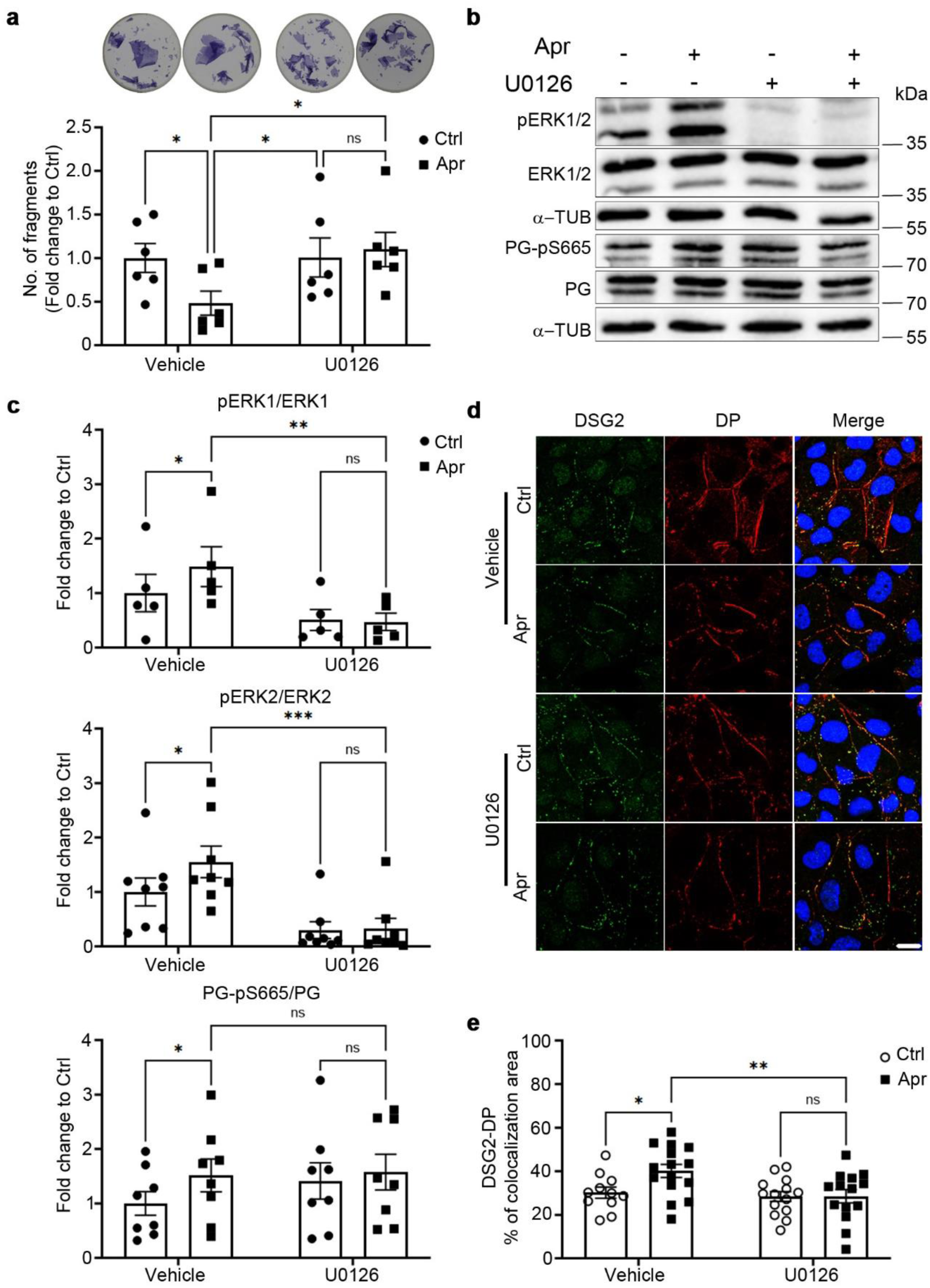
Apremilast-enhanced cardiomyocyte cohesion depends on ERK1/2 phosphorylation. **a.** Dissociation assays in HL-1 cells treated with Apr with or without the MEK inhibitor U0126 (1 µM) for 1 h. U0126 was added 30 min prior to Apr. N=7. The panel above displays representative wells for each condition **b.** Representative Western blots for PG-pS665, PG, pERK1/2, and ERK1/2 in HL-1 cells treated with Apr with or without U0126 for 1 h. α-tubulin was used as a loading control. **c.** Quantification of Western blots from **b.** N=5-8. **d.** Immunostaining and confocal microscopy were performed for DP (green) and DSG2 (red) in HL-1 cells treated with Apr with or without U0126 for 1 h. Scale bar 10 µm. **e.** Quantification of DS2-DP colocalization area. Each data point in the graph represents one confocal image performed across 3-4 experimental repeats. α-tubulin was used as a loading control. All bar graphs in the figure represent mean±SEM. Two-way ANOVA with Holm-Šidák post-hoc analysis was performed to determine the statistical significance. *p<0.05, **p<0.005 and ***p<0.0005.

### Isolation and characterization of hiPSCs from an ACM patient (ACM-hiPSCs)

So far, the results have established that apremilast enhanced cardiomyocyte cohesion in murine cardiomyocytes and cardiac slices. To further investigate whether apremilast will induce similar effects in human ACM patients, we established hiPSCs from a 14-year-old female ACM patient after she died of SCD (NP0151-11F, index patient) and her mother (NP0147-4, healthy-relative). The index patient had a biventricular phenotype and confirmed ARVC in the family according to the Task Force Criteria of 2010 ^34^. The patient carried a heterozygous pathogenic variant in *DSP c.2854G>T,* leading to a truncation of desmoplakin protein at p.Glu952*. Histological analysis of the post-mortem cardiac tissue of this ACM patient using H&E and Azan stainings revealed loss of cardiomyocytes, adipogenesis, and fibrosis of the cardiac tissue (Fig. 4a). Further, we performed immunostaining to analyze the proteins in the ICD. Analysis of desmosome proteins DP, DSG2, PG and the gap junction protein Cx43 in the post-mortem tissues revealed normal localization of the tested proteins at ICD except for DP, for which we could not detect any staining (Fig. 4b).

**Figure 4:**
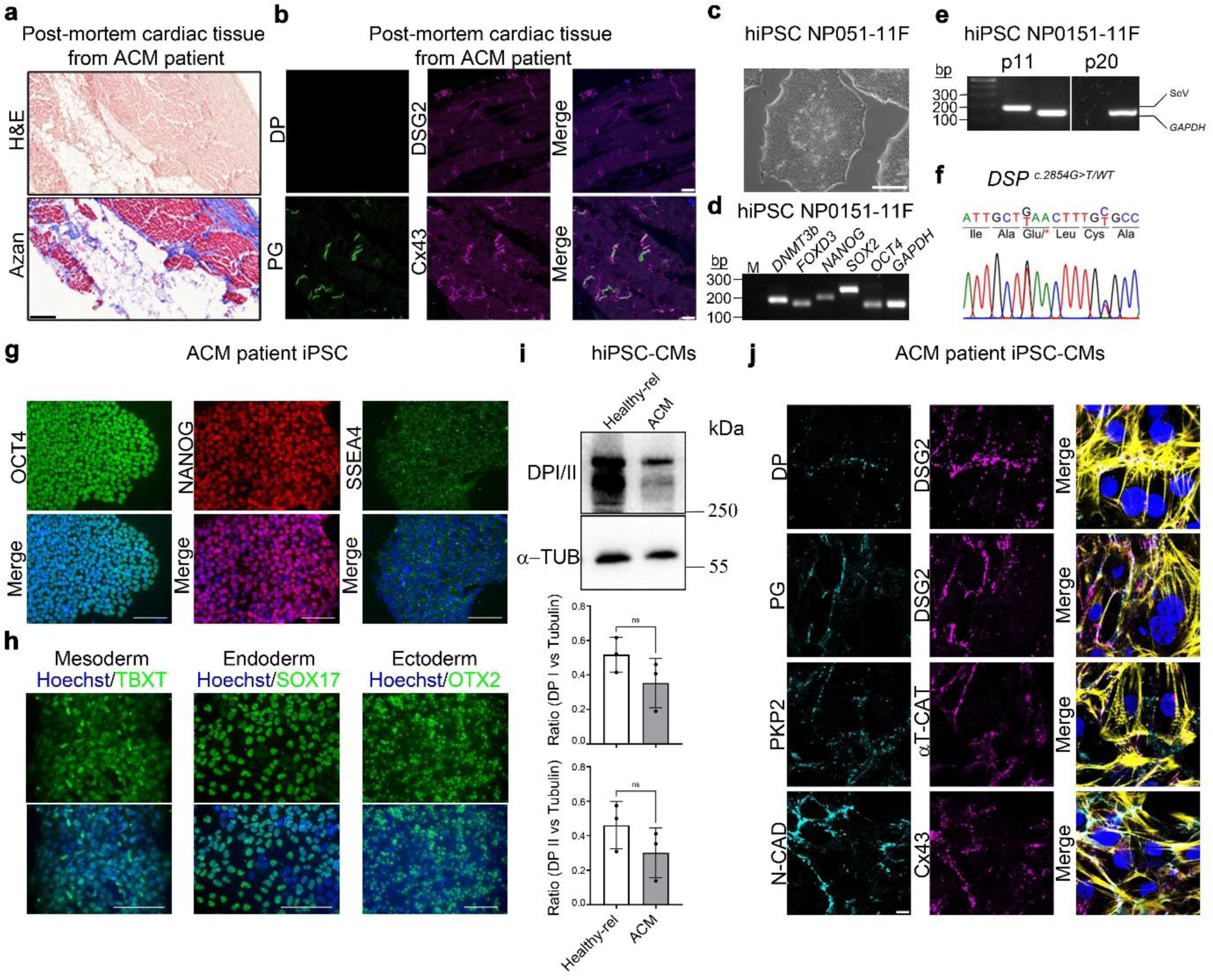
Characterization of hiPSC line NP0151-11F and hiPSC-CMs from ACM patient carrying the heterozygous mutation c.2854G>T in the *DSP* gene. The hiPSC line NP0151-11F was derived from human vein fibroblasts (hVFs) of an ACM patient carrying heterozygous mutation c.2854G>T in the desmoplakin (*DSP*) gene. **a**. Post-mortem cardiac tissue (left ventricle) of ACM patient NP0151 showing large areas of cardiomyocyte loss, adipogenesis and fibrosis. Scale bar 1 mm. **b.** Immunostaining for desmosomal proteins (DP, PG and DSG2) and gap junction protein Cx43 in the post-mortem cardiac tissue of ACM index patient revealed the absence of DSP at the ICD. Scale bar 10 µm. **c.** Colony morphology of NP0151-11F (DSP^p.Glu952*/WT^) hiPSC line. Scale bar 100 μm. **d**. Expression of indicated pluripotency-associated transcripts in ACM-hiPSC line NP0151-11F as determined by semiquantitative RT-PCR. *GAPDH* was amplified as a positive control. **e.** Confirmation of the absence of the reprogramming Sendai virus vector in NP0151-11F hiPSC master banks (passage 20) using RT-PCR. Early passage NP0151-11 hiPSCs (passage 11), which still had the reprogramming vector, were used as a positive control for the SeV presence. **f.** DNA sequencing revealed the presence of the *DSP* mutation c.2854G>T in a patient-specific hiPSC line NP0151-11F. In addition, silent heterozygous variant c.2862C>T in *DSP* was detected in NP0151-11F hiPSCs. **g.** Immunocytochemical detection of pluripotency markers OCT4, NANOG and SSEA4 in ACM-hiPSC line NP0151-11F. Scale bars 100 μm. **h.** Tri-lineage differentiation potential of the ACM-hiPSC line NP0151-11F as shown by immunocytochemical staining for TBXT (mesoderm), SOX17 (endoderm), and OTX2 (ectoderm). Scale bars 100 µm. **i.** Representative Western blots for DP I and II expression and its quantification normalized to α-tubulin in cardiomyocytes derived from healthy hiPSC line NP0040-8 and ACM patient-specific hiPSC line NP0151-11F. An unpaired student *t-test* was used to determine the statistical significance. **j.** Immunostaining for α-T-catenin (α-T-cat), DP, DSG2, PG, PKP2, N-CAD and Cx43 in cardiomyocytes derived from the ACM-hiPSC-line NP0151-11F. Actin was stained with Phalloidin Alexa Fluor 488 to show the sacromere regions of cardiomyocytes and shown in yellow in the merged image Scale bar 10 µm. Nuclei in panels **g** and **h** were counterstained with Hoechst 33342 and in panel **b** and **j** with DAPI.

The pluripotency of this hiPSC lines was confirmed by demonstrating that these cells possess typical hiPSC colony morphology (Fig. 4c and Suppl. Fig. 3a), express the selected hiPSC markers at the gene (Fig. 4d and Suppl. Fig. 3b) and protein level (Fig. 4g and Suppl. Fig. 3e), and have the ability to differentiate into derivatives of all three germ layers (Fig. 4h and Suppl. Fig. 3f), including cardiomyocytes (Suppl. Video 1 ). The hiPSC-CMs exhibited cross-striations typical for cardiac muscle cells (Suppl. Fig. 3g), and expressed all analyzed desmosomal proteins, adherens junction protein N-CAD, and the gap junction protein Cx43 (Fig. 4j). Using Sendai virus-specific PCR, we showed that the hiPSC lines NP0151-11F and NP0147-4 are free of the Sendai virus reprogramming vectors (Fig. 4e and Suppl. Fig. 3c), and the whole genome SNP profiling showed that these hiPSCs are genetically intact and retain the patient’s original genetic background (Suppl. Fig. 4). In addition, genomic DNA sequencing confirmed the presence of the pathogenic variant c.2854G>T in the *DSP* gene in this hiPSC line (Fig. 4f). Confirmation of the hiPSC line identity with the donor tissue was confirmed using Short Tandem Repeat (STR) genotyping using 14 markers, which revealed similar genetic relationship between NP0147-4 (mother) and NP0151-11F (daughter) hiPSC-lines (Suppl. Fig. 3h). In addition, DNA was isolated from fibroblasts of the ACM index-patient and from PBMCs of the healthy-relative (mother). Subsequently, the trio whole exome sequencing (WES) confirmed the pathogenic variant *c.2854G>T* in *DSP* (NM_001008844.3) in the index patient. A paternal inherited heterozygous variant of unknown significance was additionally identified in Laminin Subunit Alpha 4 (*LAMA4)*. Moreover, two polymorphisms could be detected in *CDH2* and *CCR5* (Tab. 1). Western blotting revealed that ACM-hiPSC-CMs express lower levels of DP isoforms 1 and 2 than the healthy-relative-hiPSC-CMs. However, this difference did not reach statistical significance (Fig. 4i). Immunostaining in ACM-hiPSC-CMs showed normal localization of desmosomal proteins DSG2, PKP2, PG, along with adherens junction protein N-CAD and gap junction protein Cx43. In contrast, DP staining was very faint and discontinuous along the ICD compared to DSG2 (Fig. 4j).

**Table 1:** Clinical and genetic findings in the ACM patient.

|  | **155469 (Index)** | **151514 (Mother)** |
| --- | --- | --- |
| **Clinical findings (TFC)** | 2 major criteria: pathogenic variant in *DSP*, fibrous replacement of the myocardium by autopsy (died from SCD at the age of 14 years) | Not affected |
| **Main genetic finding** |  |  |
| Gene | *DSP* | n.a. |
| Reference sequence | NM_001008844.3 |  |
| Variant | c.2854G>T, p.(Glu952Ter) |  |
| Status | heterozygous |  |
| MAF | 0 |  |
| Variant interpretation | pathogenic |  |
| **Possible genetic disease modifier** |  |  |
| Gene | *LAMA* | n.a. |
| Reference sequence | NM_001105206.3 |  |
| Variant | c.2917C>G, p.(Leu973Val) |  |
| Status | heterozygous |  |
| MAF (gnomAD) | 0.000003982 |  |
| Gene | *CCR5* | *CCR5* |
| Reference sequence | NM_000579.3 | NM_000579.3 |
| Variant | c.-301+246A>G | c.-301+246A>G |
| Status | homozygous | homozygous |
| MAF (gnomAD) | 0.4800 | 0.4800 |
| Gene | *CDH2* | n.a. |
| Reference sequence | NM_001308176.2 |  |
| Variant | c.62T>A, p.(Leu21Ter) |  |
| Status | heterozygous |  |
| MAF | 0.2007 |  |
| Comment | The variant is not present in the MANE transcript |  |

MAF: minor allele frequency, SCD: sudden cardiac death, TFC: task force criteria,

### Apremilast alleviates loss of cardiomyocyte cohesion in ACM-hiPSC-CMs

As ACM is a disease of the desmosome and desmosome formation is essential for mechanically strengthening cardiac tissue by cardiomyocyte cohesion, we questioned whether cardiomyocyte cohesion is altered in ACM-hiPSC-CMs. We performed dispase-based dissociation assays to answer this question and compared the cardiomyocyte cohesion of healthy non-relative- and healthy relative-hiPSC-CMs with ACM-hiPSC-CMs. Upon applying similar mechanical forces, ACM-hiPSC-CMs showed an increased number of fragments compared to the healthy non-relative and healthy relative-derived cardiomyocytes. This loss of cohesion was enhanced by increasing the mechanical force applied for 5 minutes (Fig. 5a and Suppl. Fig. 5a).

**Figure 5:**
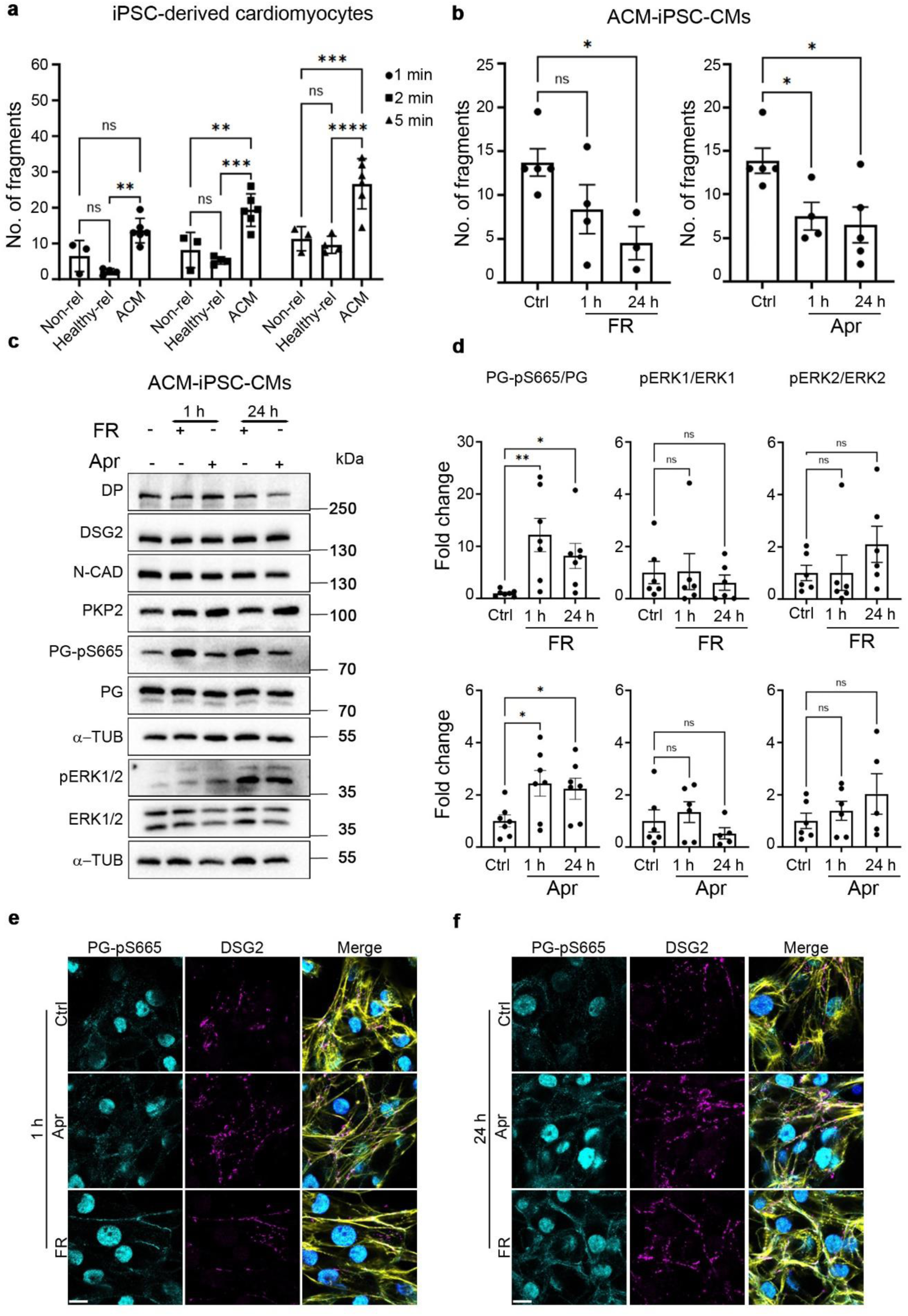
Apremilast enhances cohesion of ACM-hiPSC-CMs. **a.** Dissociation assays were performed in non-relative healthy (Non-rel), healthy mother (Healthy-rel) hiPSC- and ACM-hiPSC-derived cardiomyocytes for 1, 2, and 5 min to compare the cohesive strengths between healthy and ACM patient-derived cardiomyocytes. N=3-6. **b.** Dissociation assays were performed in ACM-hiPSC-CMs treated with FR and Apr for 1 and 24 h. N=4-5. **c.** Representative Western blots for desmosomal proteins DP, DSG2, PKP2, PG-pS665, PG, the adherens junction protein N-CAD, and the signaling molecule pERK1/2 and ERK1/2 in ACM-hiPSC-CMs treated with FR and Apr for 1 and 24 h. α-tubulin was used as a loading control. **d.** Quantification of Western blots from **c.** N=5-7. **e.** and **f.** Immunostaining and representative confocal microscopy images of PG-pS665 (cyan) and DSG2 (magenta) in ACM-hiPSC-CMs treated with FR and Apr for 1 and 24 h. Merge images show actin staining using Phalloidin Alexa Fluor 488 (yellow) and nuclei staining using DAPI (blue). All bar graphs in the figure represent mean±SEM. Scale bar 10 µm. Two-way ANOVA (**a**) and one-way ANOVA (**b** and **d**) with Holm-Šidák post-hoc analysis was performed to calculate the statistical significance. *p<0.05, **p<0.005, ***p<0.0005 and ****p<0.00005.

After establishing that ACM-hiPSC-CMs exhibited loss of cell cohesion, we investigated whether enhancing intracellular cAMP would stabilize cardiomyocyte cohesion, as previously established in murine cell lines and cardiac slices ^24–27^. Therefore, we treated ACM-hiPSC-CMs with FR and apremilast for 1 h and 24 h. Dissociation assays revealed that both FR and apremilast increased cardiomyocyte cohesion within 1 h to levels comparable to healthy hiPSC-CMs, which was stable even after 24 h (Fig. 5b compared to 5a, and Suppl. Fig. 5b). In contrast, treatment of healthy relative-hiPSC-CMs (with enhanced cohesion compared to ACM-hiPSC-CMs) revealed that apremilast enhances cardiomyocyte cohesion only after 24 h (Suppl. Fig. 5c and d). Western blot analysis of ACM-hiPSC-CMs lysates obtained after treatment with either FR or apremilast revealed increased PG-pS665 with varying intensities but no changes in ERK1/2 phosphorylation (Fig. 5c and d). FR stimulation led to a more robust PG phosphorylation than apremilast stimulation. Desmosomal proteins DP, DSG2, PKP2 and the adherens junction protein N-CAD were unaltered (Fig. 5c and Suppl. Fig. 6). Immunostaining revealed that both apremilast and FR treatment for 1 h and 24 h enhanced PG-pS665 and DSG2 at the cell junctions (Fig. 5e and f).

### Apremilast reduces arrhythmia in ex vivo murine and human ACM models

ACM patients suffer from arrhythmias, which can cause SCD. Therefore, we further investigated whether apremilast, in addition to the rescue of cardiomyocyte cohesion, can reduce arrhythmias. We measured the standard deviation of R-R intervals (SDNN) for heart rate variability as a measure for arrhythmias. First, we performed MEA analysis using *ex vivo* murine cardiac slices from *Jup^+/+^* and *Jup^-/-^* mice. MEA analysis revealed a strong increase in SDNN of *Jup^-/-^* mice cardiac slices compared to *Jup^+/+^,* which was not statistically significant due to the high variability of arrhythmia in *Jup^-/-^* tissue. Treatment with apremilast significantly decreased the SDNN in *Jup^-/-^* mice cardiac slices (Fig. 6a). To investigate the effects of apremilast in intact hearts, we performed Langendorff perfusion experiments from *Jup^-/-^* and *Jup^+/+^* mice. Apremilast treatment did not alter heart rate in both *Jup^+/+^* and *Jup^-/-^* mice (Fig. 6b and c), but ECG analysis revealed a high SDNN variability in *Jup^-/-^* hearts, which was reduced significantly after 10 min of apremilast treatment (Fig. 6d and e), which confirmed that apremilast effectively reduces arrhythmias.

**Figure 6:**
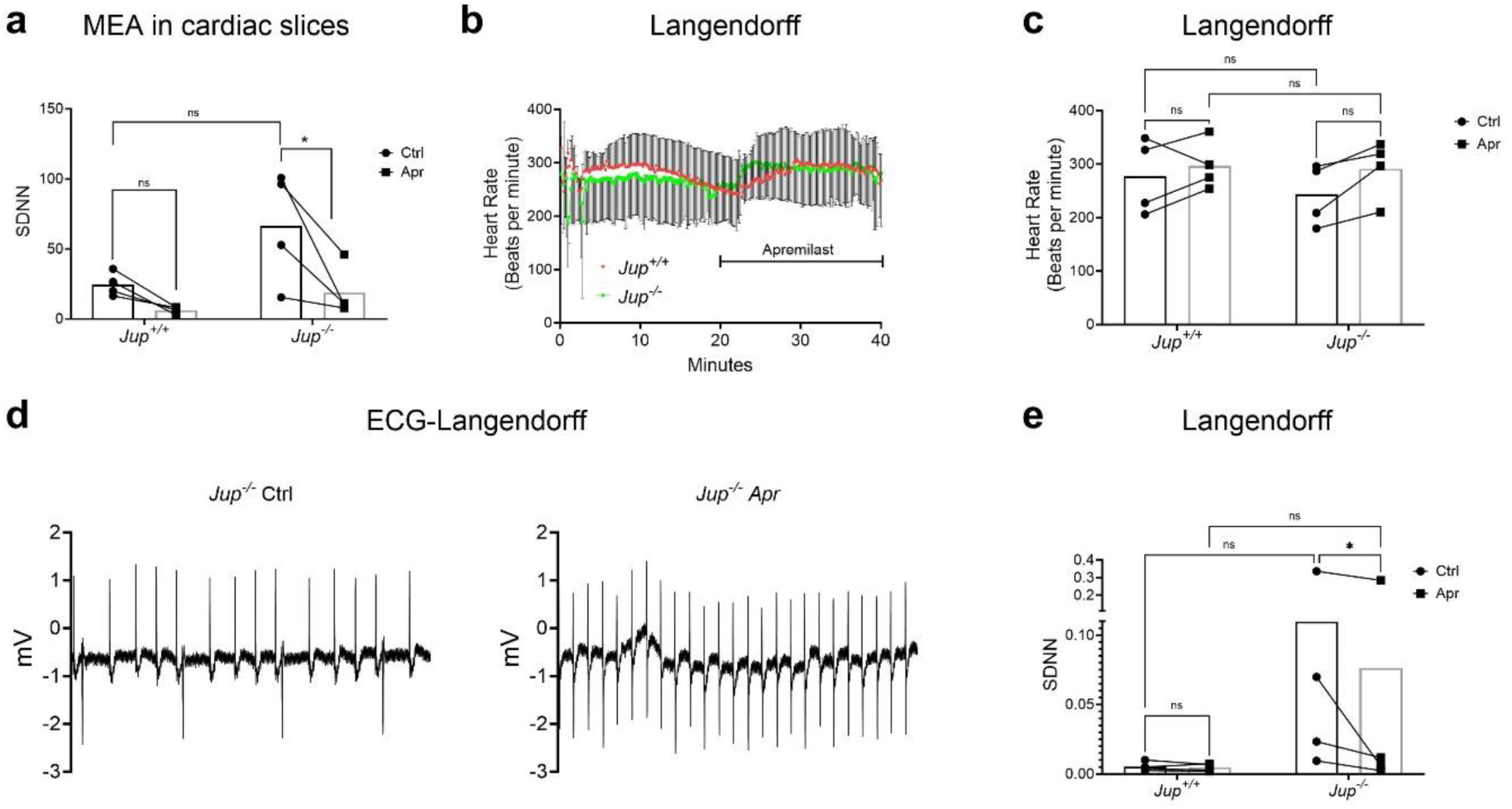
Apremilast reduces arrhythmia in a murine ACM model. Heart rate variability was measured as the standard deviation of R-R intervals (SDNN), which was used to measure arrhythmia. For this purpose, **a.** MEA was performed in the ventricular cardiac slices from *Jup^+/+^* and *Jup^-/-^* mice with or without Apr. Each data point represents one cardiac slice measured from a heart. Graph shows before-after effects of Apr, which is depicted with a connecting line between 2 data points measured on the same cardiac slice. N=3 mice per genotype. **b.** Langendorff heart perfusion experiments were performed in isolated hearts from *Jup^+/+^*and *Jup^-/-^* mice. Hearts were perfused for 20 min and then treated with Apr for 20 min. The graph shows the average heart rate measured across the whole experimental time with its corresponding SEM. N=4 mice per genotype. **c.** The graph shows the average heart rate measured in **b** for 1 min before adding Apr (Ctrl) and after 10 min of Apr treatment. **d.** Representative ECG curves (for 5 sec) obtained from the Langendorff experiments in *Jup^-/-^* mice before (upper panel) and after 10 min of Apr (lower panel). N=4. **e.** SDNN changes measured using the Langendorff perfusion experiments in the hearts of *Jup^+/+^*and *Jup^-/-^* Bar graphs in **a, c,** and **e** represent before-after plots, and graphs in **b** represent mean±SEM. Two-way ANOVA with Holm-Šidák post-hoc analysis was performed to find the statistical significance in **a, c, e** and **f**. *p<0.05, **p<0.005 and ***p<0.0005.

## Discussion

Treatment options for ACM patients exist mainly to relieve symptoms such as arrhythmia, heart failure and to prevent SCD. In most cases, β-blockers are used to control arrhythmia or ventricular tachycardia ^40, 41^. However, in some patients, beta-blockers are either ineffective ^42, 43^ or their use is limited due to side effects ^44, 45^, which then requires invasive arrhythmic procedures. The implantation of an Implantable Cardioverter Defibrillator is reserved for patients with high risk of SCD. Therefore, the current treatment strategies urge new therapeutic options for ACM patients. In contrast to the β-blockers used in ACM patients, PDE inhibitors, which act by enhancing the ionotropic state of the heart, are very effective in treating heart failure ^46, 47^. Moreover, PDE inhibitors were also shown to be effective in treating inflammation ^48, 49^, which could be a possibility to treat patients with hot phases or myocarditis which occurs in *DSP* cardiomyopathy. In line with this, the PDE-4 inhibitor apremilast was applied effectively in patients suffering from the desmosomal autoimmune skin disease pemphigus vulgaris ^36, 37^. Further, a recent study showed that apremilast mitigated the loss of keratinocyte cohesion induced by pemphigus autoantibodies^35^. In addition, recent evidence suggest that stabilization of cell adhesion could be beneficial in treating desmosome-related diseases^50^. Based on the above observations, we hypothesized that apremilast can enhance the integrity and mechanical stability of desmosomes at cell junctions, enhance cardiomyocyte cohesion, and reduce arrhythmia in a murine ACM model.

We have previously reported that increasing intracellular cAMP levels and modulating other signaling mechanisms lead to strengthening of cardiomyocyte cohesion, primarily in murine models such as the HL-1 atrial cell line and murine cardiac slice cultures ^24–27, 39^. One significant challenge hindering progress in the study of ACM is the limited access to human tissues and suitable human cells for research. While animal models have provided valuable insights, an alternative approach involves hiPSCs, which can be differentiated into CMs, offering a robust in vitro model for studying ACM ^16, 18, 51–54^. The ACM-hiPSC cell line NP0151-11F was established from fibroblasts obtained from tissue of the deceased ACM patient carrying the point mutation c.2854G>T in the *DSP* gene that results in a premature stop codon and a truncated protein product at p.Glu952*. WES confirmed the presence of the pathogenic variant c.2854G>T in the *DSP* gene in the human ACM-hiPSC-CMs derived from this hiPSC line. The premature termination of the protein sequence at p.Glu952* affects DP interaction with desmin, whereas interaction domains for PG and PKP2 remained intact ^55–57^. Furthermore, WES also revealed genetic variants of unknown significance in *LAMA4* and two polymorphisms in *CDH2* and *CCR5* genes in this patient. However, a CCR5 gene variant was also found in the healthy mother.

After the establishment of ACM-hiPSCs and their differentiation into CMs, a dissociation assay was implemented to measure cardiomyocyte cohesion, comparing it to healthy non-relative and healthy mother hiPSC-CMs. For the first time, we demonstrate here that ACM-hiPSC-CMs exhibit impaired cohesion. Notably, treatment with either apremilast or FR restored the loss of cohesion within 24 h comparable to healthy-relative cardiomyocyte levels. These observations were further corroborated in a murine ACM model showing that both apremilast and FR enhanced cardiomyocyte cohesion in cardiac slice cultures from *Pkp2^-/-^* mice. In line with this, apremilast and FR enhanced baseline cell adhesion in WT cardiac slices and cultured HL-1 cardiomyocytes, which was in agreement with previous studies ^24–27, 58^. The observations that cardiomyocyte cohesion is impaired in human iPSC-derived cardiomyocytes carrying a pathogenic *DSP* variant and that adhesion can be restored by pharmacological interventions similar to intact tissue cultures from a murine model of ACM with *Pkp2* deletion indicate that strengthening the desmosomal integrity and mechanical stability might represent a new therapeutic approach for ACM patients. ACM is a disease with complex pathogenicity where factors other than genetic variants contribute to disease phenotype and progression, including inflammation during so-called ‘hot phases’ as well as physical exercise. The observation that cardiomyocyte cohesion is impaired in cultured cells from patients in whom ACM is caused by a pathogenic variant of a desmosomal component such as *DSP* in the absence of additional stimuli indicates that loss of cell adhesion is a primary event in ACM pathogenesis similar to other desmosome diseases such as pemphigus ^8, 29^ and that restoration of cell adhesion may be a primary goal of pharmacological therapy in ACM. Thus, increasing cAMP by apremilast or inhibiting EGFR by erlotinib might be promising new treatment options for ACM patients since they were proven to be effective in enhancing desmosome integrity and, thereby, cell adhesion in different experimental models of ACM and pemphigus, as shown in this study and previous reports ^35, 38, 59^.

We also investigated the molecular mechanism of how apremilast restored cell cohesion in ACM-hiPSC-CMs and enhanced cell adhesion in HL-1 cells and murine slice cultures. Apremilast induced translocation of DSG2, PG and DP to cell contacts and phosphorylation of PG at serine 665 in all models except *Jup^-/-^*mice. Increased intracellular cAMP leads to PG-pS665 and enhances cardiomyocyte cohesion mirrors observations in murine cell lines and cardiac slices and is in agreement with the effects of FR as reported before ^8, 24–27^. These data show that the mechanism of positive adhesiotropy is present in human cardiomyocytes. However, since the increase of cAMP induced by apremilast was ten times less than that observed after FR treatment, whereas the effect on cell adhesion was comparable, it can be concluded that higher cAMP levels, which may cause side effects, are not required to strengthen cardiomyocyte adhesion.

Because we established previously that increased intracellular cAMP enhances cardiomyocyte cohesion via PKA-mediated PG-pS665 ^24, 26, 27, 60^, we tested this phenomenon utilizing heart slice cultures from PG serine 665 phospho-deficient (JUP-S665A) mice. Interestingly, apremilast enhanced cardiomyocyte cohesion in the absence of PG serine phosphorylation. Further, apremilast enhanced DSG2 and PG at the ICD, suggesting that apremilast can enhance cardiomyocyte cohesion by compensatory effects in a model where PG phosphorylation is genetically blunted. However, in the absence of PG in heart slice cultures from *Jup^-/-^* mice, the effects of apremilast were abolished altogether, revealing that PG is strictly necessary for apremilast-mediated effects on cardiomyocyte cohesion.

We described previously that ERK1/2 activation plays a crucial role in cAMP-mediated cardiomyocyte cohesion in murine cardiomyocytes and heart slice cultures ^25^. Therefore, we tested this phenomenon utilizing HL-1 cells. Apremilast, similar to FR in our previous study ^25^, caused activation of ERK1 and required ERK1/2 for its positive adhesiotropic effect. In contrast, neither apremilast nor FR activated ERK1/2 in ACM-hiPSC-CMs, suggesting that the underlying mechanisms enhancing desmosomal adhesion may be at least in part different in human and murine cardiomyocytes. This should be considered when studying ACM pathogenesis in different experimental models, and it would suggest that human iPSC-CMs have advantages over murine models.

Most ACM patients suffer from arrhythmias, and in some, arrhythmias can lead to SCD ^5, 9, 61^. Therefore, we next investigated whether apremilast could stabilize the rhythmicity of spontaneous beating in different cardiac models. MEA analysis and Langendorff perfusion of intact murine hearts and cardiac slice cultures from the PG-deficient ACM model and the SDNN of R peaks was used as a measure for arrhythmia. The experiments revealed a significant reduction of arrhythmia after apremilast treatment. This supports previous findings that a molecular approach to enhance desmosomal adhesion can reduce arrhythmia, as shown before using a DSG2-linking peptide ^62^.

In summary, we demonstrate here for the first time that cell adhesion was impaired in cardiomyocytes derived from an ACM patient and that apremilast effectively restored desmosome-mediated cardiomyocyte cohesion and reduced arrhythmia in ACM model. Encouragingly, apremilast was proven to inhibit doxorubicin-induced apoptosis and inflammation in the heart ^48^ and was also used as an anti-inflammatory drug in autoimmune and inflammation-related diseases such as psoriasis, arthritis and inflammatory bowel disease ^63–65^. Recently, autoimmunity and inflammation were observed as prominent features in ACM^21, 23, 51, 66^ and therapeutic modulation of inflammation in ACM hearts revealed an effective new mechanism-based therapy for ACM ^51^. Further, clinical data from a large patient population revealed no adverse risks of major cardiac events in psoriatic arthritis patients ^67^. Overall, the studies mentioned above, combined with our data in the present study, support apremilast as a promising new therapeutic option for ACM.

## Materials and methods

### Reagents

The supplementary file details all the important reagents, mediators, primers, and antibodies used in this study and the missing methods in the main manuscript.

### HL-1 cell culture

As described in our previous studies, the murine atrial cardiac myocyte cell line HL-1 was maintained in Claycomb medium containing norepinephrine ^23, 25, 26^. Cells seeded for experiments were incubated in Claycomb medium without norepinephrine to avoid basal adrenergic stimulation.

### Dissociation assays in HL-1 cells

For dissociation assays, experiments were performed as explained previously ^23, 25, 26^ and in supplementary methods.

### Murine models

All mice lines used in this study were generated and bred as described previously for *JupS665A*^35^*, Jup^-/-^* ^24–26, 38, 39, 62^ and *Pkp2*^−/−^ ^38^. For experiments age- and sex-matched littermates, 8-14-week-old mice, were used. Euthanasia was performed using isofluran (Iso-Vet, 1000 mg/g) in an E-Z anesthesia system (#EZ-SA800-OS, World precision instruments, Germany). Mice were briefly anesthetized with 5% isoflurane until the mouse became immobile, mostly within 1 to 2 minutes. A toe pinch was applied to ensure that the animal was anesthetized, then cervical dislocation was performed, and hearts were excised for further analysis. All the animal experiments confined to the principles outlined in the Declaration of Helsinki. Ethics statement was provided in the section ‘Study approval’.

### Sex as a biological variable

Both male and female mice were included, and similar findings were found in both sexes.

### Murine cardiac slice culture and dissociation assay from mice

Murine cardiac slice cultures, dissociation assays, and lysates for Western blots were obtained exactly as published previously ^38^. Details of this can be found in supplementary methods.

### Human tissue collection for hiPSC generation

Fibroblasts were isolated at the Institute of Human Genetics of the LMU Munich from the vein of the index patient (ID-number NP0151) collected after the autopsy at the Pathology Department of the Munich Clinic Schwabing. The human dermal fibroblasts (hDFs) used to generate the NP0040-8 hiPSC line were isolated from a full-thickness skin sample obtained aseptically from a healthy male subject by punch biopsy. The tissue specimens were placed into cell culture dishes, cut into small pieces, and cultured in the Dulbecco’s modified Eagle medium (DMEM) supplemented with 10% fetal bovine serum (FBS), 2% Glutamax, 1% non-essential amino acids (NEAA), 100 U/ml penicillin, 100 μg/ml streptomycin, and 100 µM β-mercaptoethanol (β-ME) until confluent outgrowing fibroblasts appeared. The human vein fibroblasts (hVFs) were then collected by dissociation with TrypLE Express (Thermo Fisher Scientific, Waltham, MA, USA), expanded for an additional 2-3 passages and cryopreserved in aliquots for future use.

The NP0147-4 hiPSC line was generated from peripheral blood mononuclear cells (PBMCs) that were obtained from the index patient’s healthy mother. Whole blood (∼30 ml) was collected by venepuncture into BD Vacutainer CPT vials and PBMCs were immediately isolated by centrifugation at RT following the manufacturer’s protocol. PBMC aliquots were frozen in 10% DMSO in liquid nitrogen for future use.

### Generation of hiPSCs

To generate insertion-free hiPSCs, cryopreserved hVFs or hDFs (collectively referred to as hFs) were thawed and passaged in hF medium for approximately two passages before reprogramming was initiated by transduction with Sendai virus (SeV) vectors included in the CytoTune-iPS 2.0 Sendai Reprogramming Kit (Thermo Fisher Scientific) at the recommended multiplicity of infection (MOI). The transduced hFs were cultured in hF medium without antibiotics at 37 °C at 5% CO_2_ under normoxic conditions for one day, and the medium was replaced with fresh hF medium. The cells were then cultured in hF medium until day 7, changing the medium every other day. On day 7, cells were dissociated with 0.05% trypsin/EDTA, plated at densities of 2, 5 and 10 x10^4^ cells per well of a 6-well plate coated with 0.5 μg/cm^2^ vitronectin (VTN-N, Thermo Fisher Scientific) and cultured in hF medium for a further 2 days. On day 8, the hF medium was replaced with E8 medium containing 100 ng/ml basic fibroblast growth factor (bFGF, Peprotech, Hamburg, Germany), and cells were cultured with a medium change every two days until pluripotent stem cell-like colonies appeared, usually within 15-28 days after reprogramming. Initial hiPSC colonies were manually dissected and small cell clusters were transferred to fresh vitronectin-coated 6- or 12-well plates. Each harvested colony was cultured separately in complete E8 medium and after several passages, those clones that showed typical hiPSC morphology with no or low level of spontaneous differentiation were selected for further expansion, characterization and cryopreservation.

### Cultivation and cryopreservation of hiPSCs

hiPSCs were regularly passaged by Versene (Thermo Fisher Scientific) upon reaching about 70-80% confluency at the split ratio of 1:4 to 1:6 and grown in complete E8 medium supplemented with 10 μM ROCK inhibitor Y27632 (Selleck Chemicals, Cologne, Germany) for the first 24 h. All hiPSC cultures were regularly checked for mycoplasma contamination using the MycoAlert® Mycoplasma Detection Kit (Lonza, Cologne, Germany). Master banks were prepared after hiPSC lines were confirmed to be SeV vector-free. Passage numbers at vector clearance varied between individual clones and usually occurred between passage numbers 7 and 19. Cells were banked as small cell clusters after dissociation with Versene in E8 medium supplemented with 10% DMSO and stored in liquid nitrogen until further use.

### Maintenance of hiPSC lines

hiPSCs were maintained in Essential 8^TM^ Medium (Gibco^TM^, cat #A1517001) on Vitronectin (Gibco^TM^, cat #A14700)-coated 6-well plates (Greiner, cat #657160) for NP0040-8 control hiPSC line or on Matrigel^TM^ (Corning, cat #734-1440)-coated 6-well plates for NP0147-4 and NP0151-11F hiPSC lines in a humidified incubator at 37 °C with 5% CO_2_. Cells were passaged when 70-80% confluent (once every 3-5 days) by washing twice with DPBS without Mg^2+^ and Ca^2+^ (Gibco^TM^, cat #14190250) and incubating with Versene (Gibco^TM^, cat #15040066) for 3 to 5 min at RT. After removing Versene cell clusters were gently washed from the well with Essential 8^TM^ Medium (for NP151-11F hiPSCs with 2 µM ROCK inhibitor) and transferred to a newly coated 6-well plate at split ratios between 1:6 to 1:20.

### Differentiation of hiPSC lines

hiPSCs were differentiated into cardiomyocytes by using small molecules to modulate Wnt signaling pathways. When maintained cultures reached 80% confluency, hiPSCs were seeded as single cells by washing twice with DPBS without Mg^2+^ and Ca^2+^ and incubating with TrypLE^TM^ Express (Gibco^TM^, cat #12604021) with 10 µl DNase I (2500 U/ml, Thermo Scientific^TM^, cat #90083) per 1 ml TrypLE^TM^ for 5 min at 37°C. The reaction was stopped with Essential 8^TM^ Medium supplemented with 10 µM ROCK inhibitor (Y27632, Adooq, cat # A11001-5) and cells were filtered through a 40 µm cell strainer (corning®, cat #431750). After centrifugation and resuspension into fresh Essential 8^TM^ Medium supplemented with 10 µM ROCK inhibitor, 0.6 to 0.8 x 10^6^ single cells were seeded into each well of a 6-well plate. Cells were maintained in Essential 8^TM^ Medium and differentiated when cells reached 90% confluency. First, cells were treated for exactly 24 h with 8 µM CHIR99021 (Sigma-Aldrich, #SML1046) in RPMI GlutaMAX^TM^ (Gibco^TM^, cat #61870010) with 50 µg/ml ascorbic-acid (WAKO Chemicals Europe, cat # 013-12061), B27 supplement without insulin (Thermo Scientific^TM^, cat #A1895601) and penicillin/streptomycin (Gibco, cat # 15140-122). Cells were recovered in RPMI-B27minus insulin for 48 h before treated with 5 µM XAV939 (Sigma Aldrich, cat #X3004-5MG) and 5 µM IWP2 (Tocris, cat #3533/10) at day 3 for 48 h. At day 5 medium was changed to RPMI-B27minus insulin and from day 7, when beating monolayers started forming, medium was changed to fresh RPMI-B27plus insulin and refreshed every 2 days. Cells were replated for experiments between day 8 and 12.

### RT-PCR

The clearance of the reprogramming SeV vectors in established hiPSC lines and expression of the pluripotency-associated markers OCT3/4, NANOG, SOX2, FOXD3 and DNMT3b at the transcript level was determined using semi-quantitative RT-PCR. Primers used for these analyses are listed in the Supplementary Table 1. For monitoring the vector clearance, total RNA was isolated using TRIzol (Thermo Fisher Scientific) from uninfected cells (negative control), from cells freshly transduced with SeV vectors (positive control), and from established hiPSC clones at different stages of expansion. To determine the expression of the pluripotency-associated markers, RNA samples were isolated using the same method from an aliquot of banked cells. RNA was reverse transcribed into cDNA using random hexamers for priming and SuperScript II Reverse Transcriptase (Thermo Fisher Scientific) following the manufacturer’s recommendations. Negative control in RT reactions included all reaction components except RTase. Target sequences were amplified by PCR using DreamTaq Green PCR Master Mix (2X) (Thermo Fisher Scientific) using the following cycling program: initial denaturation for 90 s at 95 °C followed by 35 cycles of 30 s denaturation at 95 °C, 30 s annealing as specified in the Supplementary Table 1, 30 s extension at 72 °C, and 5 min final extension at 72 °C. GAPDH PCR-product was used as a housekeeping gene reference. PCR products were analyzed by agarose gel electrophoresis and visualized using ethidium bromide.

### Mutation analysis

To confirm the presence of the disease-causing pathogenic variant c.2854G>T in *DSP*, the genomic DNA was isolated from hiPSC lines using the DNeasy Blood & Tissue Kit (Qiagen) according to the manufacturer’s recommendations. The gene segment harbouring the mutation was amplified by PCR using the primers listed in Supplementary Table 1, the PCR product was cleaned up using the QIAquick PCR purification Kit (Qiagen), and the eluted DNA samples were sent to Eurofins Genomics (Ebersberg, Germany) for sequencing using the forward and the reverse PCR primer.

### Exome analysis

Sample preparation and bioinformatic analysis of the index patient NP0151-11F and her mother [healthy-relative (NP147-4)] were performed at the Institute of Human Genetics at the Technical University of Munich. DNA was isolated from fibroblasts (patient) or blood lymphocytes (mother), respectively. Exome sequencing was performed using the Twist Human Exome 2.0 Plus Comprehensive Exome Spike-in and Mitochondrial Panel. Sequencing was performed on the NovaSeq 6000 as a 100 bp paired-end run (Illumina, San Diego, California). Primary and secondary bioinformatic analysis were performed using the variant analysis pipeline EVAdb (https://github.com/mri-ihg/EVAdb). In particular, the Burrows-Wheeler Aligner (BWA) was used to map the generated data to the human genome sequence GRCh37. Single nucleotide variants (SNVs) and insertions/deletions (indels) were determined using SAMtools, PINDEL and the Genome Analysis Toolkit (GATK) package. Copy number variations (CNVs) were identified using ExomeDepth.

Exome-derived variant calls were annotated based on genomic coordinates, effect on gene product, and minor allele frequency within population control resources (gnomAD and in-house control exome dataset). Variants were further annotated for deleteriousness by calculating predictions from CADD, SIFT, and PolyPhen. To establish genetic causality and prioritize candidate genes, we considered various online repositories and in silico metrics such as ClinVar, HGMD, PubMed, OMIM, Uniprot Genotype-Tissue GTEx, Human Protein Atlas and gnomAD probability of being loss-of-function intolerant (pLI) and missense z-scores. After annotation, the variants were filtered using lists of sequencing artifacts, and calls with suboptimal quality parameters were eliminated. We applied a recessive filter for homozygous and compound heterozygous variants with a minor allele frequency (MAF) < 1%. CNVs and mtDNA variants with a MAF filter < 0.01% or < 1% were evaluated. We applied a filter for variants that were listed as “pathogenic” or “likely pathogenic” in the ClinVar database without an additional MAF filter. Additionally, we explicitly filtered for variants in the *CDH2* gene without any MAF filter since our group previously showed that the autoantibodies in ACM patients cleaved and reduced N-CAD binding ^23^. The Integrative Genomics Viewer (IGV) was used to inspect the filtered SNVs, indels, and CNVs visually.

### Dissociation assays with hiPSC-CMs

Dissociation assay experiments were performed similarly to that of HL-1 cells with some modifications. hiPSC-CMs were seeded between days 8 and 12 of differentiation and plated with 2×10^5^ cells per cm^2^ density on Matrigel™-coated plates. hiPSC-CMs were grown until day 20 while changing medium every 2 days. hiPSC-CMs were treated with respective mediators for 1 h or 24 h. Subsequently, the cells were washed with HBSS, treated with Liberase-DH (0.065 U/ml, Sigma-Aldrich #5401054001) and Dispase II (2.5 U/ml, Sigma-Aldrich, #D4693-1G), and incubated at 37 °C until the cell monolayer detached from the wells (usually within 10-15 min). Then, the enzyme mix was carefully removed and replaced by HBSS. Mechanical stress was applied by horizontal rotation at 1250 rpm for 1 to 10 min. The number of fragments was determined utilizing a binocular stereomicroscope (Leica Microsystems, Wetzlar, Germany) and ImageJ analysis software.

### Western blotting

After respective treatments, HL-1 cells, hiPSC-CMs or cardiac slices were lysed in SDS lysis buffer supplemented with protease and phosphatase inhibitors (cOmplete Protease Inhibitor Cocktail, Roche; CO-RO and PhosStop, Roche, PHOSS-RO). SDS-polyacrylamide gel electrophoresis was performed using 10-15 µg of protein for HL-1 cells or hiPSC-CMs and 40 µg for cardiac slices. Further details are provided in Supplementary materials. The membranes were incubated with antibodies of interest in 5% BSA/TBST or Roti-Block overnight at 4 °C (BSA: VWR; cat # 422361V). Further details of primary antibody and secondary antibody concentrations are provided in Supplementary Table 2. Membranes were washed 3 times in TBST for 5 min and incubated in respective goat anti-mouse/rabbit-HRP conjugated secondary antibody at 1:20000 dilution in TBST for 1 h at room temperature. After incubation, the membranes were washed 3 x for 10 min in TBST and immunoreactive bands were detected with ECL solution Super Signal West (Thermo Scientific; cat # 34577) after exposure in a FluorChemE system (ProteinSimple California, USA).

### Immunostaining of HL-1 cells, hiPSC-CMs and murine cardiac slices

Immunostaining in HL-1 cardiomyocytes and murine cardiac slices was performed as described previously ^25, 38^ and explained further in supplementary file. Detailed information on immunostainings in hiPSC-CMs are provided in the supplementary file.

### Multielectrode array (MEA) analysis in murine cardiac slices

MEA analysis in murine cardiac slices was performed as explained before ^58, 62^ with some modifications. MEA2100-60-System equipped with 60MEA200/30iR-Ti-gr electrode chambers (both Multichannel Systems) was used for this analysis. 300-μm-thick cardiac slices were cut using Vibratome as indicated in the supplementary methods and were washed with HBSS buffer and transferred to Claycomb medium without norepinephrine supplemented with 10% fetal bovine serum and 10 μg/ml penicillin and streptomycin and incubated at 37 °C, 5% CO_2_, for 15 min. Then cardiac slices were placed on MEA electrodes containing 500 μl of cardiac slice medium, and measurements were performed with MC_Rack software 4.6.2 (Multichannel Systems). Slices from either *Jup^+/+^* or *Jup^-/-^* hearts were analyzed by MEA for 5 min (Control) before the addition of 1 μM apremilast. Slices were then incubated at 37 °C and 5% CO_2_ for 30 min and further measured for 5 min. SDNN was measured as the standard deviation of the beat-to-beat intervals.

### Langendorff heart perfusion

Adult murine hearts (from 12-14 weeks old mice) were used for Langendorff heart perfusion experiments using a commercially available apparatus (AD Instruments), and the results were analyzed using LabChart8 software as described previously ^39^. Mice were sacrificed by cervical dislocation under isoflurane anesthesia and underwent subsequent preparation of the hearts in less than 5 min. Hearts were perfused in the retrograde Langendorff mode with heparin-free carbogen-gassed Krebs-Henseleit buffer supplemented with 18.8 nM norepinephrine at 37 °C with 60 mm H_2_O constant pressure. ECG was recorded via needle electrodes. After recording the baseline for 20 min, 1 µM apremilast was added to the perfusion solution for the indicated time using a perfusion pump. Heartbeats and SDNN intervals were measured using LabChart8 software.

### Statistics

Statistical analyses were performed using GraphPad Prism 8 or higher version. Two-tailed Student’s *t* tests or 1- or 2-way ANOVA with post hoc tests were applied after outlier removal and are explained in figure legends. Data are represented as mean ± SEM. Significance was assumed for *P* ≤ 0.05.

### Study approval

Animal handling was done under the guidelines from Directive 2010/63/EU of the European Parliament and approved by the regional government of Upper Bavaria (Gz. ROB-55.2-2532.Vet_02-19-172, for *Jup* mice) or the local ethics committee of the government of Lower Franconia (RUF-55.2.2-2532-2-663 and 55.2.22-2532-2-955 for *Pkp2* mice).

The Ethics Committee of the Medical Faculty of the University of Cologne (authorization number 14-306) approved patient tissue collection and use for generating hiPSCs. Further hiPSC study at the LMU was approved by the Ethics Committee approval at LMU (23-0203).

## Data availability

The data corresponding to this manuscript can be obtained from the corresponding authors upon considerable request.

## Author contributions

KS, OD, JK, CW, SP, MS, SOF, SM, JO, NLCSW and SY acquired the data. KS, OD, TŠ and SY analyzed the data. EG handled the biosample, sequencing platform, and bioinformatics regarding the trio WES. DSW analyzed the trio WES data. RB provided the materials for hiPSCs and cardiac tissue. TW and BG provided hearts from *Pkp2*^−/−^ mice. hiPSCs were generated, and protocols to differentiate hiPSCs into cardiomyocytes were established in the lab of TŠ. SY drafted the manuscript. SY, TŠ and JY made critical revisions to the manuscript for important intellectual content and SY and JY designed the research. All authors proofread the manuscript.

## Funding

The LMU Munich supported this work through the Funding program for research and teaching (FöFoLe) to SY and JW and the Deutsche Forschungsgemeinschaft grant WA2474/11-1 and WA2474/14-1 to JW. hiPSC lines were generated and characterized with the support of the grants from the Innovative Medicines Initiative of the EU and EFPIA (Agreement number IMI JU–115582) and the Köln-Fortune Program to T.Š.

## Acknowledgements

We thank Kathleen Plietz and Kilian Skowranek (Institute of Anatomy, Faculty of Medicine, Ludwig-Maximilian-University (LMU) Munich), Rebecca Dieterich and Maike Kreutzenberg (Center for Physiology and Pathophysiology, University Hospital Cologne and Medical Faculty, University of Cologne, Germany) for technical assistance and Stefan Herms (Life & Brain GmbH, Department of Genomics, Bonn, Germany) for SNP genotyping and bioinformatics analysis.

## Notes

Conflict of interest: The authors have declared that no conflict of interest exists.

### Competing Interest Statement

The authors have declared no competing interest.

