## Supplementary material for "Apremilast improves cardiomyocyte cohesion and arrhythmia in different models for arrhythmogenic cardiomyopathy": Suppl. file

### Supplementary materials and methods

#### cAMP assay

The intracellular concentration of cAMP was measured by cAMP Enzyme Immunoassay Kit (Sigma-Aldrich, #CA200-1KT). After the respective treatments, cells were washed, and 0.1 M HCl was added to the respective well, which was further processed according to the manufacturer's protocol. Absorbance was measured at a wavelength of 450 nm using the TECAN Infinite 200 PRO microplate reader (Tecan Deutschland GmbH).

#### Dissociation assays in HL-1 cells

Cells were seeded at 125000 cells per $cm^2$ on cell culture plates coated with 0.02% gelatin and 25 µg $ml^{-1}$ fibronectin, grown for 72 h, and then washed with HBSS and treated with Liberase-DH (0.065 U $ml^{-1}$, Sigma-Aldrich Munich, #5401054001) and Dispase II (2.5 U $ml^{-1}$, Sigma-Aldrich, #D4693-1G). The cells were then incubated at 37 °C until the cell monolayer detached from the wells. Then, the enzyme mix was carefully removed from the wells and replaced by HBSS. Mechanical stress was applied by horizontal rotation on an orbital shaker (Stuart SSM5 orbital shaker) at 1.31 g for 10-15 min. The number of fragments were determined by counting utilizing a binocular stereomicroscope (Leica Microsystems, Wetzlar, Germany).

#### Murine cardiac slice culture

After anesthetizing mice with isoflurane, they were sacrificed by cervical dislocation, and the hearts were excised and put into pre-cooled slicing buffer, embedded in low-melt agarose, and sliced with a vibratome (LeicaVT1200S vibratome) into 200 µm sections for dispase, immunostaining and Western blots, and 300 µm sections for MEA. For treatments with drugs, consecutive slices were used to minimize slice-to-slice variability. After respective treatments, slices were processed for lysate preparation, dissociation assays, and immunostainings. For immunostainings, slices were embedded in NEG-50 frozen section medium and stored at either -20 °C or -80 °C until further processing.

#### Dissociation assays in murine cardiac slices

Twelve weeks old age- and sex-matched littermates of wild-type and genetically modified mice were used for experiments. Mice were sacrificed by cervical dislocation, hearts were immediately placed in pre-cooled oxygenated cardiac slicing buffer, embedded in low melt agarose, and 200 µm thick slices were cut with a LeicaVT1200S vibratome (Leica Biosystems). For dissociation assays, slices were washed gently with HBSS, transferred to pre-warmed cardiac slices medium in separate wells of a 12-well plate, and incubated for 1 hour with indicated compounds at 37 °C, 5% $CO_2$. Then, Liberase-DH and Dispase II were added together for 30 min. Next, MTT (3-(4, 5-dimethylthiazolyl-2)-2, 5-diphenyltetrazolium bromide) was added to check for cell viability and mechanical stress was applied using an electronic pipette with 1 ml pipette tip by pipetting up and down for 10-20 times. However, the same number of pipetting were always applied between the control and treated cardiac slices of a mouse. Finally, the same mechanical stress was applied to the other slices of the same experiment. Then the content of each well was filtered using a 70 µm nylon membrane, which resulted in mostly single cells or couple of cells as clusters and pictures of the well were taken and stitched together using the AutoStich software. Only the number of dissociated single cardiomyocytes stained with MTT were counted for quantification. The number of dissociated cells was normalized to the respective consecutive untreated control slice of the same mouse to reduce slice-to-slice variability in slice size.

**Murine cardiac slice lysates**

When working with murine cardiac slice cultures, slices were treated, washed with TBS, and snap-frozen in liquid nitrogen. To prepare the tissue lysates 400 µl-500 µl of SDS lysis buffer supplemented with protease and phosphatase inhibitors was added and the samples were transferred into gentleMACS M-tubes (Miltenyi Biotec, 130-093-236) and dissolved using the protein_01_01 program of the gentleMACS OctoDissociator for 1 min (Miltenyi Biotec, 130-095-937) and processed according to the manufacturer protocol. Isolated protein samples were collected in 1.5 ml Eppendorf tubes and stored at -20 °C for short-term storage or at -80 °C for long-term storage. Western blots were performed as explained in material and methods.

**Immunostaining of HL-1 cells, hiPSC-CMs and murine cardiac slices**

HL-1 cells were seeded on coverslips and grown for 3 days before respective treatments were performed. hiPSC-CMs were seeded between days 8 and 12 of differentiation, plated with a density of 2x105 cells per $cm^2$ on Matrigel™-coated plates, and grown until day 20. After respective treatments, cells were fixed with either 2% (HL-1 cells) or 4% (hiPSC-CMs) paraformaldehyde (PFA). Sections of murine cardiac slices (7 µm thick) were cut with a cryostat (HM500OMV, Microm); transferred onto glass slides and adhered by warming up to 38 °C. After fixation with 2% PFA, the samples were permeabilized with 0.1% Triton X-100 and blocked with BSA and normal goat serum (BSA/NGS). Primary antibodies diluted in PBS were added and incubated overnight at 4 °C in a wet chamber. Primary and secondary antibody details were provided in Supplementary Table 2. When working with HL-1 and hiPSC-CMs, PBS was used for washing steps, while, for murine tissue samples, 50 mM $NH_4Cl$ in PBS was used. After washing, species-matched, fluorophore-coupled secondary antibodies were added for 60 min at room temperature. In the case of wheat germ agglutinin (WGA; Thermo Fisher Scientific, W11261), the dye was added together with the secondary antibodies and DAPI, at a concentration of 2 µg/ml. After washing, coverslips were mounted on microscope slides using ProLong Diamond Antifade Mountant (Thermoscientific, cat # P36961). Slides were analyzed using a Leica SP5 II confocal microscope (Leica) with a 63× oil objective and a numerical aperture of 1.4 using Immersol 518F (Zeiss). Images were acquired at room temperature with the LAS-AF software. Colocalization analyses were performed as described before (25, 38). In brief, the membrane region was marked as a region of interest; in this region, the amount of stained pixels in the other channel, where colocalization was to be quantified, was measured, and a colocalization ratio at the membrane was calculated. Measurement of staining width in immunostained murine cardiac slices, was performed as described previously (38).

**Trilineage differentiation potential of iPSCs**

The potential of iPSCs to differentiate into the three germ layers (mesoderm, endoderm, and ectoderm) was evaluated using the STEMdiff™ Trilineage Differentiation Kit (Stem Cell Technologies, Cologne, Germany) following the manufacturer's protocol. The expression of the germ-layer-specific markers was assessed by immunocytochemistry using antibodies against Brachyury (T) for mesoderm (Cell Signaling Technology, cat # 81694, 1:400), Sox17 for endoderm (Cell Signaling Technology, cat # 81778, 1:800), and OTX2 for ectoderm (Santa Cruz Biotechnology, cat # sc-514195, 1:200). Cells were fixed with 4% PFA on day 5 for mesoderm and endoderm lineages and day 7 for the ectoderm lineage. Staining with antibodies and microscopy were performed as described in supplementary materials.

**Genetic analysis of iPSCs**

Karyotypic integrity and genetic identity of iPSCs was assessed by a global single nucleotide polymorphism (SNP) analysis using the Human Omni2.5 Exome-8 BeadChip v1.3 from Illumina (San Diego, CA, USA). To this end, the genomic DNA was isolated from an aliquot of banked iPSCs and matched somatic cells of the corresponding patient using the DNeasy Blood & Tissue Kit (Qiagen, Hilden, Germany) according to the manufacturer's instructions. DNA was prepared for hybridization to bead chip arrays according to the manufacturer's protocol. The raw intensity data (*.idat files) were analyzed by the Genotyping Module of Illumina Genome Studio software version 2.0.3. Normalized Log R Ratio and B Allele Frequency for all the available probes in each sample were extracted.

**Immunostaining for pluripotency markers**

Immunocytochemistry was used to assess the expression of the pluripotency-associated markers in hiPSC lines and their potential to differentiate to derivatives of all three germ layers. For this purpose, undifferentiated hiPSCs or their differentiated derivatives were fixed with 3.7% paraformaldehyde (PFA) for 10 min, permeabilized with PBS containing 0.25% Triton X-100 and 0.5 M $NH_4Cl$, and blocked for 60 min with 1x Roti-Block (Roth, Karlsruhe, Germany). Samples were then stained overnight at 4 °C with primary antibodies specific for OCT3/4 (Santa Cruz Biotechnology, Heidelberg, Germany, Cat. No. sc-5279, 1:200), NANOG (Cell Signaling Technology Europe, Leiden, The Netherlands, Cat. No. 4903-S, 1:200), TRA-1-60 (BD Biosciences, Heidelberg, Germany, Cat. No. 560121, 1:100), TRA-1-81 (Santa Cruz, Cat. No. Sc-21706, 1:100) and SSEA4 (Santa Cruz Biotechnology, Cat. No. Sc-21704, 1:200). All antibodies were diluted to their working concentrations in 1x Roti-Block. Signals were visualized after staining for 1 h at room temperature with AlexaFluor 488- or AlexaFluor 555-conjugated secondary antibodies (Thermo Fisher Scientific) diluted at 1:1000 in 1x Roti-Block. Nuclei were counterstained with Hoechst 33342 (1:1000). Samples were embedded in ProLong Gold Antifade Reagent (Thermo Fisher Scientific) and observed on Axiovert Microscope (Carl-Zeiss Microimaging, Oberkochen, Germany) equipped with the image processing software Axiovision 4.5.

**Western blotting**

Each sample was mixed with Laemmli sample buffer containing DTT and boiled for 5 min at 95 °C and loaded onto a 7.5% polyacrylamide gel with stacking gel. The gel electrophoresis was run in a Bio-Rad Mini-PROTEAN® Tetra Cell system, starting with a constant 80 V for 20 min, followed by a constant 120 V for an additional 45 to 60 min. 5 µl of Page Ruler Plus (Thermo Fisher Scientific; #26619) was also run on the same gel for molecular weight determination. The separated proteins were transferred onto a 0.45 µm nitrocellulose membrane (Thermo Fisher Scientific; #LC2006) for 90 min at constant 350 mA using Bio-Rad Mini-PROTEAN® Tetra System. To block non-specific binding sites, the membranes were incubated at room temperature for 1 h in 0.1 M TBS with 0.1% Tween 20 (TBST) containing 5% non-fat dry milk (Sigma Aldrich; #70166-500G) or with 1x Roti-Block (Roth, ROTI®-Block, #A151.2).

**Supplementary Table 1.** Primer sequences used in PCR reactions.

| **Primer name** | **Sequence (5' → 3')** | **Annealing temperature (°C)** | **Expected product size (bp)** |
|---|---|---|---|
| SeV-F | GCATCACTAGGTGATATCGAGC | 55 | 181 |
| SeV-R | ACCAGACAAGAGTTTAAGAGATATGTATC | | |
| DNMT3b-F | GTCGTGCAGGCAGTAGGAAA | 60 | 175 |
| DNMT3b-R | GCCATTTGTTCTCGGCTCTG | | |
| FOXD3-F | GCAACTACTGGACCCTGGAC | 60 | 145 |
| FOXD3-R | CTGTAAGCGCCGAAGCTCT | | |
| NANOG-F | TTTGGAAGCTGCTGGGGAAG | 60 | 194 |
| NANOG-R | GATGGGAGGAGGGGAGAGGA | | |
| OCT3/4-F | CATGTGTAAGCTGCGGCCC | 66 | 268 |
| OCT3/4-R | GCCCTTCTGGCGCCGGTTAC | | |
| SOX2-F | ACGGCTGGAGCAACGGCAGC | 66 | 137 |
| SOX2-R | GGAGTTGTACTGCAGGGCGC | | |
| GAPDH-F | GCTCATGACCACAGTCCAT | 60 | 142 |
| GAPDH-R | ACCTTGCCCACAGCCTTG | | |
| DSP-F1-Seq | ACAGGCTGACTAGAAAGTAACC | 56 | 546 |
| DSP-R1-Seq | CACCTACCACCAGTCAAAGAA | | |

**Supplementary Table 2.** Antibodies and blocking/dilution reagents used for Western blotting (WB) and immunofluorescence (IF).

| Antibody | Blocking / Dilution WB | Blocking / Dilution IF |
|---|---|---|
| DP (mAb, mouse, Progen, #690003) | Roti-Block / 1:500 in Rotiblock | BSA/NGS / 1:100 in BSA/NGS |
| DSG2 (mAb, mouse, Progen, #61002) | 5% milk / 1:1000 in BSA/TBST | / |
| DSG2 (mAb, rabbit, Progen, #610121) | / | BSA/NGS / 1:100 in BSA/NGS |
| N-CAD (mAb, mouse, BD Transduction, #610921) | 5% milk / 1:1000 in BSA/TBST | BSA/NGS / 1:100 in BSA/NGS |
| PKP2 (mAb, mouse, Progen, #651167) | 5% milk / 1:25 in BSA/TBST | BSA/NGS / 1:20 in BSA/NGS |
| $pPG^{S665}$ (mAb, mouse, homemade hybridoma) | 5% milk / 1:20 in BSA/TBST | BSA/NGS / 1:20 in BSA/NGS |
| PG (mAb, mouse, Progen, #61005) | 5% milk / 1:1000 in BSA/TBST | BSA/NGS / 1:100 in BSA/NGS |
| α-Tubulin (mAb, mouse, Abcam, #ab7291) | 5% milk / 1:4000 in BSA/TBST | / |
| $pERK1/2^{T202/Y204}$ (mAb, mouse, Santa Cruz Biotechnology, #sc-7383) | 5% milk / 1:1000 in BSA/TBST | / |
| ERK1/2 (pAb, rabbit, Cell Signalling Technology, #9102) | 5% milk / 1:1000 in BSA/TBST | / |
| Cx43 (pAB, rabbit, Sigma, #SAB4501175) | / | BSA/NGS / 1:100 in BSA/NGS |
| α-T-catenin (pAb, rabbit, Proteintech, #13974-1-AP) | / | BSA/NGS / 1:100 in BSA/NGS |
| Goat-anti-mouse-HRPO (Dianova, #115-035-045) | 1:10000 in TBST | / |

| | | |
|---|---|---|
| Goat-anti-rabbit-HRPO (Dianova, #111-035-045) | 1:10000 in TBST | / |
| Goat-anti-mouse Cy3 (Dianova, # 115-165-164) | / | 1:600 in PBS |
| Goat-anti-rabbit Cy3 (Dianova, # 111-165-003) | / | 1:600 in PBS |
| Coat-anti-mouse Cy5 (Dianova, # 115-175-071) | / | 1:600 in PBS |
| Goat-anti-rabbit Cy5 (Dianova, # 111-175-144) | / | 1:600 in PBS |
| Phalloidin Alexa 488 (Molecular Probes\Life technologie, # A 12°9) | / | 1:400 in PBS |
| DAPI, Roche, #10236276001 | / | 1:2000 in PBS |
| Brachyury (TBXT) for mesoderm (Cell Signaling Technology, # 81694) | / | 1:400 |
| SOX17 for endoderm (Cell Signaling Technology, # 81778) | / | 1:800 |
| OTX2 for ectoderm (Santa Cruz Biotechnology, Cat. No. sc-514195) | / | 1:200 |
| OCT3/4 (Santa Cruz Biotechnology, # sc-5279) | / | 1:200 in Roti-Block |
| NANOG (Cell Signaling Technology # 4903-S) | / | 1:200 in Roti-Block |
| TRA-1-60 (BD Biosciences, # 560121) | / | 1:100 in Roti-Block |
| TRA-1-81 (Santa Cruz, # Sc-21706) | / | 1:100 in Roti-Block |
| SSEA4 (Santa Cruz Biotechnology, # Sc-21704). | / | 1:200 in Roti-Block |

**Supplementary figures:**

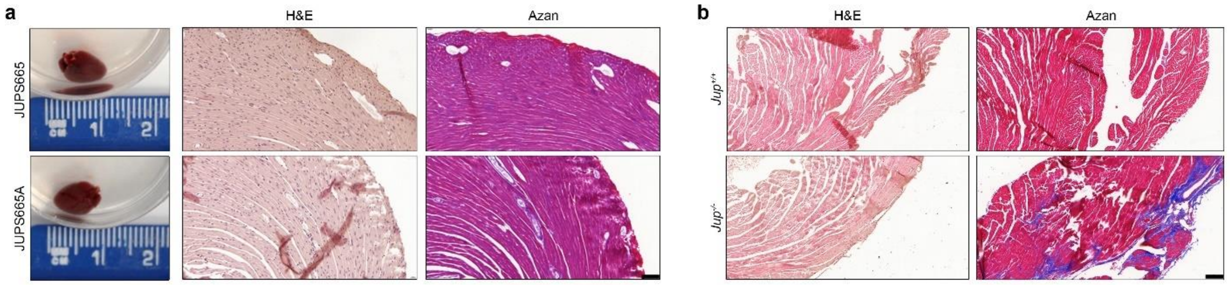

**Supplementary figure 1: Histo-morphological analysis of the hearts from Jup S665A and *Jup*$^{-/-}$ mice.**

**a.** Morphological images of hearts from JupS665 and JupS665A mice on the left panel. H & E and Azan staining revealed no gross changes in JupS665A mice compared to their wild-type littermates, JupS665. Hearts were excised from mice aged between 12-15 weeks. Scale bar 100 µm. **b.** H & E and Azan staining reveal cardiomyocyte loss and fibrosis in *Jup*$^{-/-}$ mice compared to their wild-type littermates, *Jup*$^{+/+}$ mice. Hearts were excised from 12-week-old mice. Images are representative of a minimum 3 mice observed per genotype. Scale bar 100 µm.

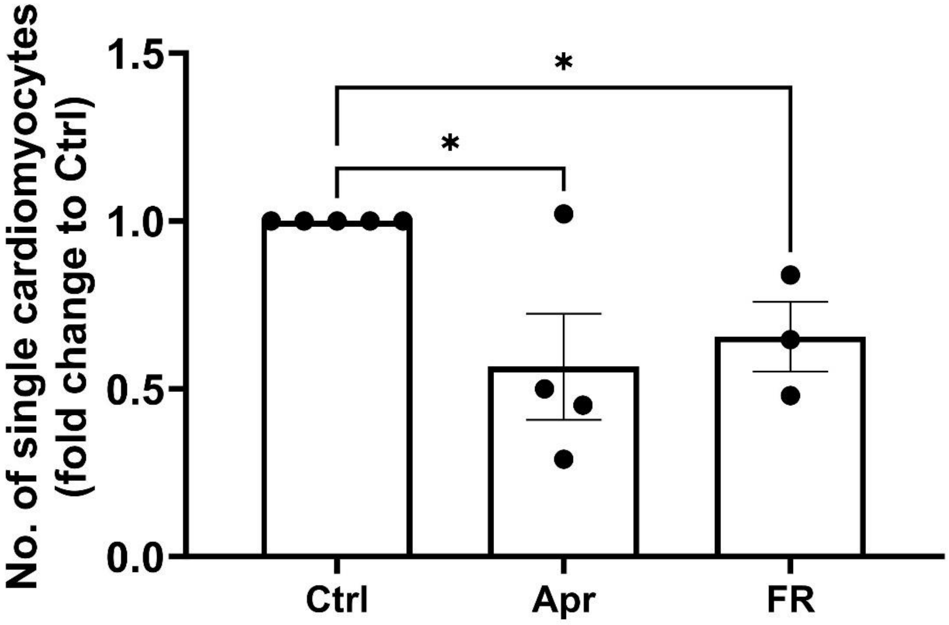

**Supplementary figure 2: Dissociation assay using cardiac slices prepared from hearts of *Pkp2*$^{-/-}$ mice.**

Dissociation assays were performed with heart slices of *Pkp2*$^{-/-}$ mice treated with Apr and FR for 1 h, N=3-4 mice per condition. One-way ANOVA with Holm-Šidák post-hoc analysis was performed to determine the statistical significance. * $p<0.05$.

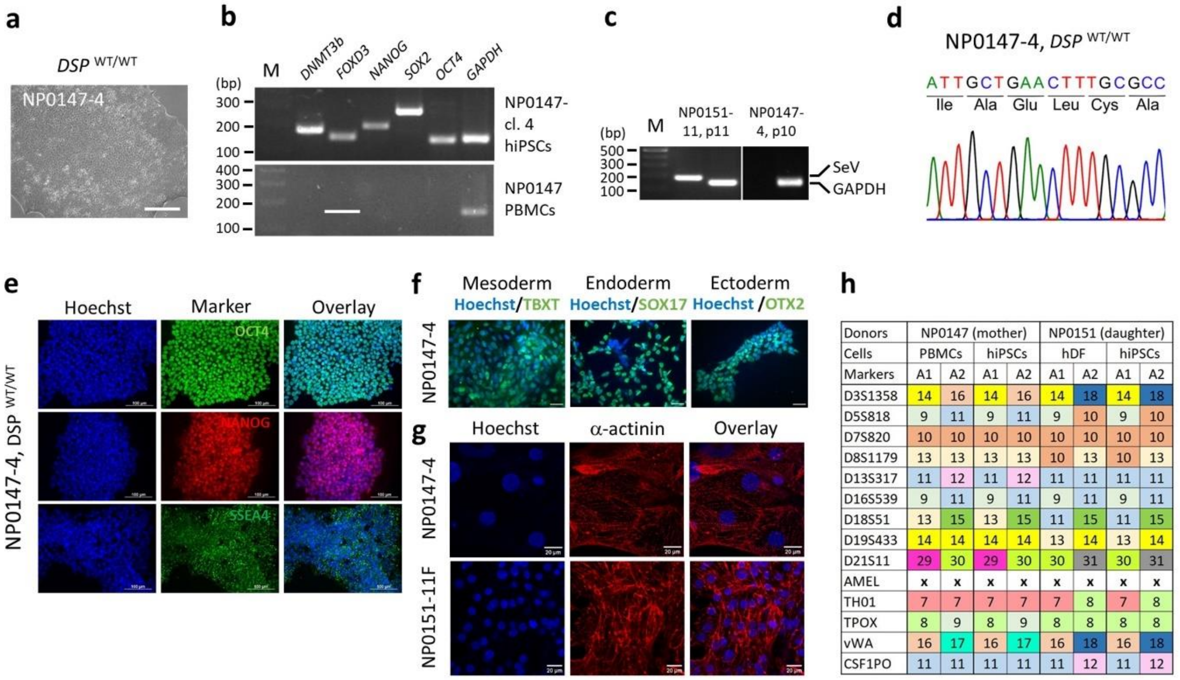

| Donors | NP0147 (mother) | | | | NP0151 (daughter) | | | |
|---|---|---|---|---|---|---|---|---|
| Cells | PBMCs | | hiPSCs | | hDF | | hiPSCs | |
| Markers | A1 | A2 | A1 | A2 | A1 | A2 | A1 | A2 |
| D3S1358 | 14 | 16 | 14 | 16 | 14 | 18 | 14 | 18 |
| D5S818 | 9 | 11 | 9 | 11 | 9 | 10 | 9 | 10 |
| D7S820 | 10 | 10 | 10 | 10 | 10 | 10 | 10 | 10 |
| D8S1179 | 13 | 13 | 13 | 13 | 10 | 13 | 10 | 13 |
| D13S317 | 11 | 12 | 11 | 12 | 11 | 11 | 11 | 11 |
| D16S539 | 9 | 11 | 9 | 11 | 9 | 11 | 9 | 11 |
| D18S51 | 13 | 15 | 13 | 15 | 11 | 15 | 11 | 15 |
| D19S433 | 14 | 14 | 14 | 14 | 13 | 14 | 13 | 14 |
| D21S11 | 29 | 30 | 29 | 30 | 30 | 31 | 30 | 31 |
| AMEL | x | x | x | x | x | x | x | x |
| TH01 | 7 | 7 | 7 | 7 | 7 | 8 | 7 | 8 |
| TPOX | 8 | 9 | 8 | 9 | 8 | 8 | 8 | 8 |
| vWA | 16 | 17 | 16 | 17 | 16 | 18 | 16 | 18 |
| CSF1PO | 11 | 11 | 11 | 11 | 11 | 12 | 11 | 12 |

**Supplementary figure 3: Characterization of hiPSC lines from the healthy-relative (NP0147-4) and ACM-patient (NP0151-11F).**

The hiPSC line NP0147-4 was generated from peripheral blood mononuclear cells (PBMCs) of the patient's mother (healthy relative). **a**. Colony morphology of a wild-type NP0147-4 (*DSP* *WT/WT*) hiPSC lines. Scale bar 100 µm. **b**. Expression of indicated pluripotency-associated transcripts as determined by semiquantitative RT-PCR. mRNA isolated from PBMCs obtained from the donor NP0147 was used as negative control and *GAPDH* was amplified as a positive control. cl – clone. **c**. Confirmation of the absence of the reprogramming Sendai virus vector in NP0147-4 hiPSC master banks using RT-PCR. Early passage NP0151-11 hiPSCs, which still had the reprogramming vector, were used as a positive control for the SeV presence. **d.** DNA sequencing in the NP0147-4 hiPSC line from the ACM patient's healthy mother **e**. Immunocytochemical detection of pluripotency markers OCT4, NANOG and SSEA4 in healthy relative hiPSC lines. Scale bars 100 µm. **f.** Tri-lineage differentiation potential of the healthy relative (NP0147-4) hiPSC lines as shown by immunocytochemical staining for TBXT (mesoderm), SOX17 (endoderm), and OTX2 (ectoderm). Scale bars 100 μm. **g.** Both NP0147-4 and NP0151-11F hiPSC lines give rise to cardiomyocytes as shown by immunocytochemical staining for cardiac $\alpha$-actinin. Scale bar 20 μm. Nuclei in panels **e**,**f**, and **h** were counterstained with Hoechst 33342. **h.** Confirmation of the hiPSC line identity with the donor tissue using Short Tandem Repeat (STR) genotyping with 14 markers. The genetic relationship between NP0147 (mother) and her daughter (donor NP0151) was also verified.

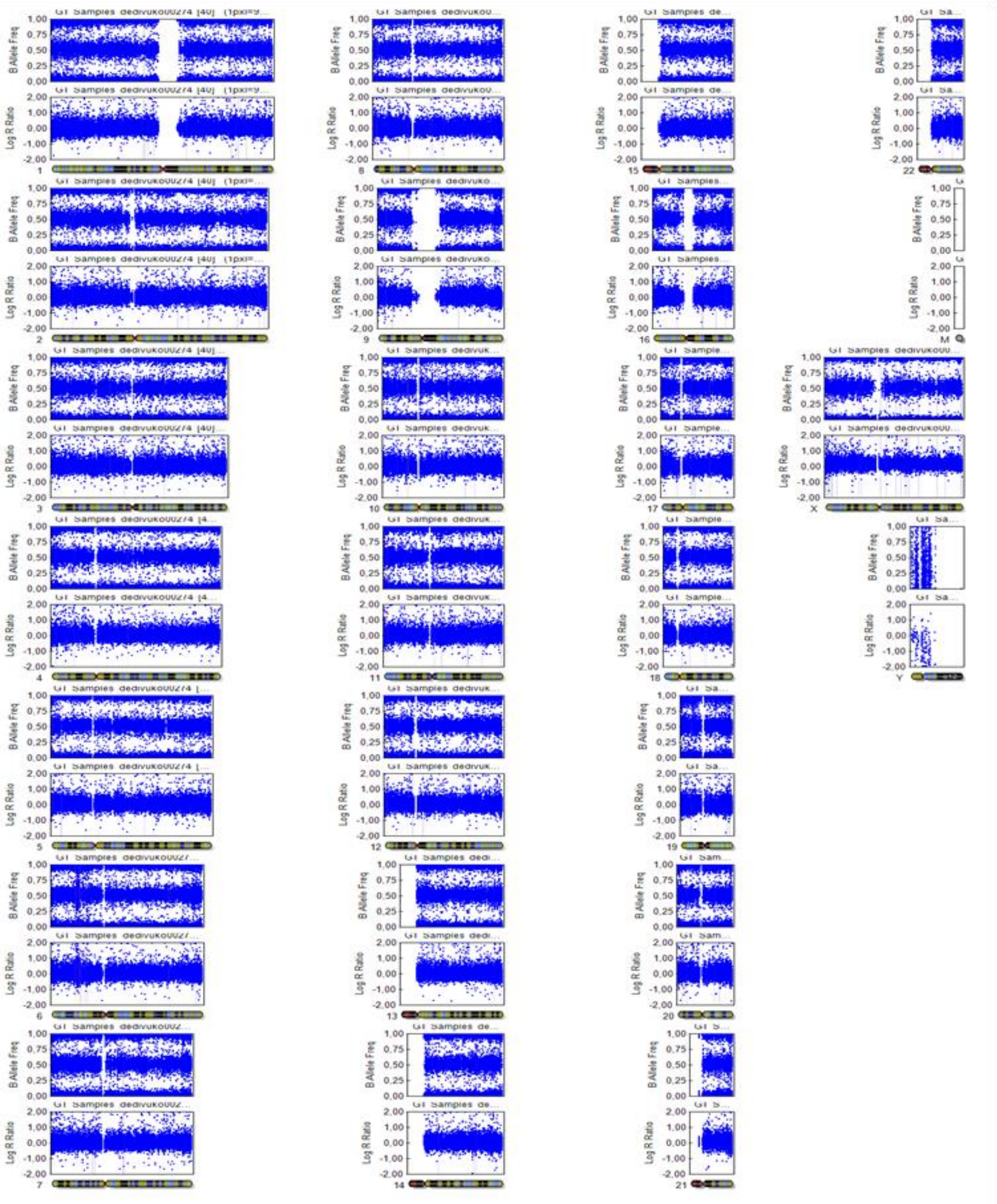

**Supplementary figure 4: Molecular karyotype of the ACM-hiPSC line NP0151-11F.**

The analysis was done using genomic DNA isolated from an aliquot of cells prepared for a Master hiPSC bank at passage 9+18. The image shows the B-allele frequencies (upper panels) and log2 R ratios (lower panels), plotted for each chromosome. Each dot represents a single nucleotide polymorphism (SNP).

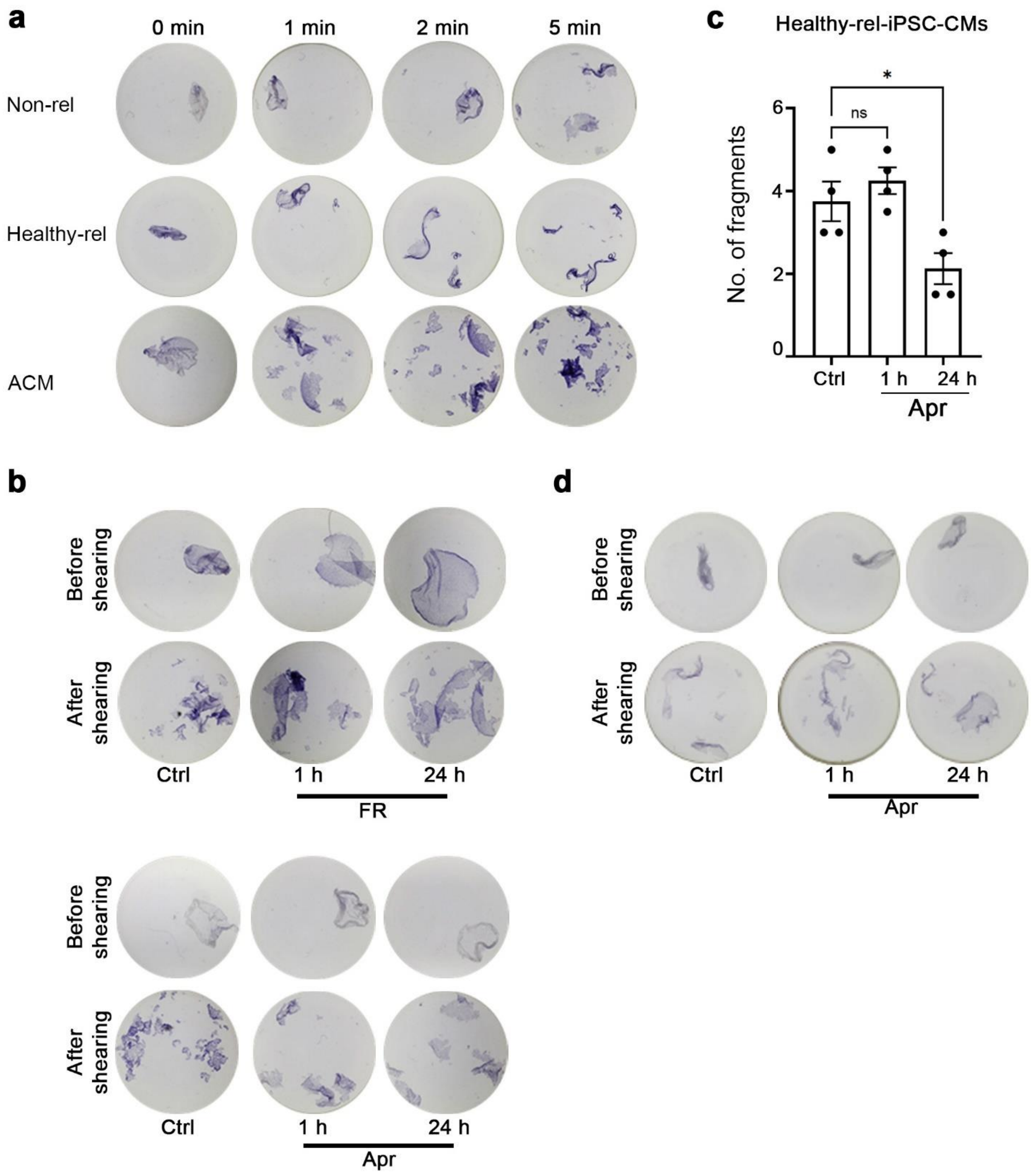

**Supplementary figure 5: Representative wells for dissociation assays performed with ACM-hiPSC-CMs and apremilast effect on healthy-rel-iPSC-CMs cohesion.**

Representative wells of dissociation assays in figure **5a** and **b** were depicted in **a.** and **b,** showing the wells before and after shear stress**. c.** Healthy relative (healthy-rel) -hiPSC-CMs were treated for 1 h and 24 h, and then dissociation assays were performed. Bar graph represents mean±SEM. One-way ANOVA with Holm-Šidák post-hoc analysis was performed to determine the statistical significance *$p<0.05$. **d.** Representative wells of dissociation assay performed in **c.**

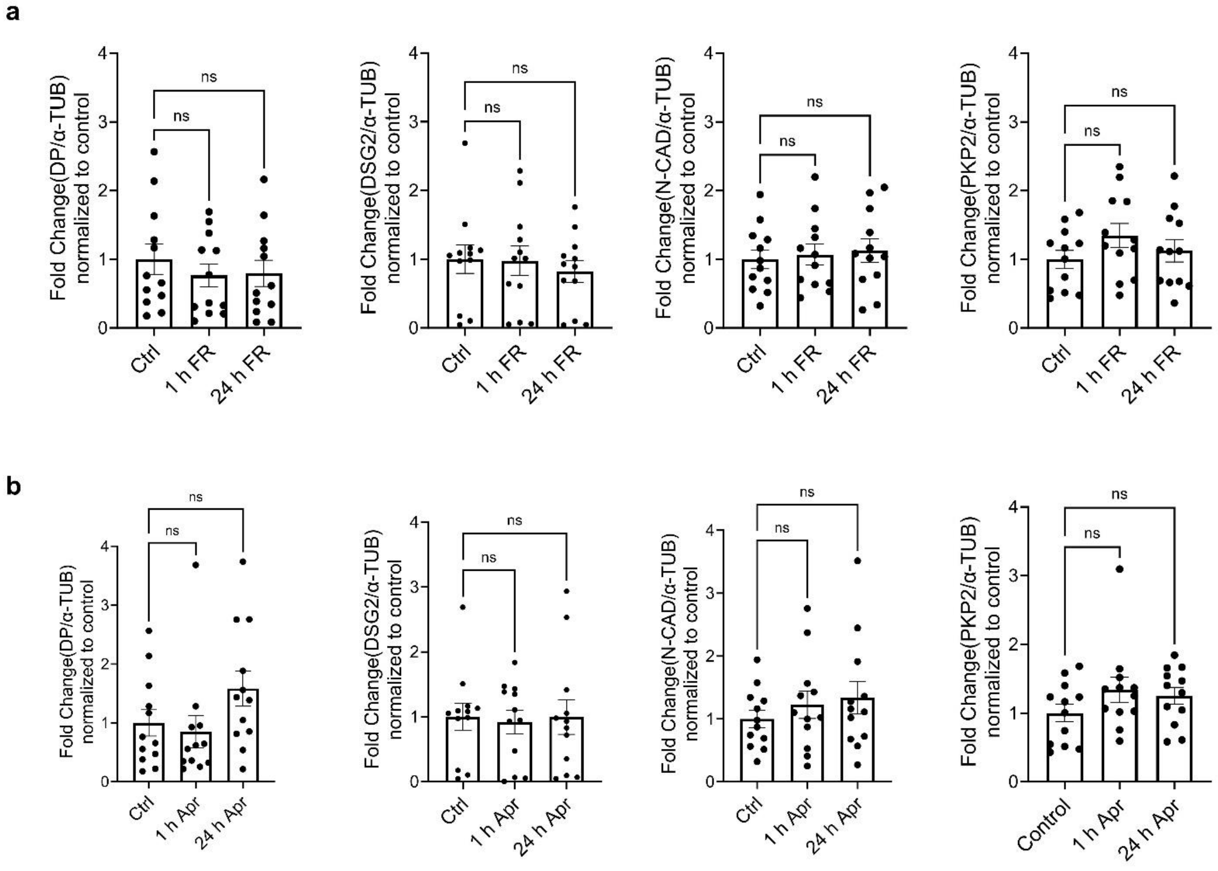

**Supplementary figure 6: Quantification of Western blots from figure 5c.**

Quantification of Western blots for desmosomal proteins DP, DSG2, PKP2 and the adherens junction protein N-CAD in ACM-hiPSC-CMs treated with **a.** FR and **b.** apremilast (Apr) for 1 h and 24 h. N=12. All bar graphs in the figure represent mean±SEM
